# DREAMER: Exploring Common Mechanisms of Adverse Drug Reactions and Disease Phenotypes through Network-Based Analysis

**DOI:** 10.1101/2024.07.20.602911

**Authors:** Farzaneh Firoozbakht, Maria Louise Elkjaer, Diane E. Handy, Rui-Sheng Wang, Zoe Chervontseva, Matthias Rarey, Joseph Loscalzo, Jan Baumbach, Olga Tsoy

## Abstract

Adverse drug reactions (ADRs) are a major concern in clinical healthcare, significantly affecting patient safety and drug development. The need for a deeper understanding of ADR mechanisms is crucial for improving drug safety profiles in drug design and drug repurposing. This study introduces DREAMER (Drug adverse REAction Mechanism ExplaineR), a novel network-based method for exploring the mechanisms underlying adverse drug reactions and disease phenotypes at a molecular level by leveraging a comprehensive knowledge graph obtained from various datasets. By considering drugs and diseases that cause similar phenotypes, and investigating their commonalities regarding their impact on specific modules of the protein-protein interaction network, DREAMER can robustly identify protein sets associated with the biological mechanisms underlying ADRs and unravel the causal relationships that contribute to the observed clinical outcomes. Applying DREAMER to 649 ADRs, we identified proteins associated with the mechanism of action for 67 ADRs across multiple organ systems, e.g., ventricular arrhythmia, metabolic acidosis, and interstitial pneumonitis. In particular, DREAMER highlights the importance of GABAergic signaling and proteins of the coagulation pathways for personality disorders and intracranial hemorrhage, respectively. We further demonstrate the application of DREAMER in drug repurposing and propose sotalol (targeting KCNH2), ranolazine (targeting SCN5A, currently under clinical trial), and diltiazem (indicated drug targeting CACNA1C and SCN3A) as candidate drugs to be repurposed for cardiac arrest. In summary, DREAMER effectively detects molecular mechanisms underlying phenotypes emphasizing the importance of network-based analyses with integrative data for enhancing drug safety and accelerating the discovery of novel therapeutic strategies.

## 1. Introduction

Adverse drug reactions (ADRs) are important concerns in pharmacology and healthcare. They are the leading cause of mortality and drug withdrawals (Tatonetti 2012). Gaining a deeper understanding of ADRs is essential for enhancing drug safety profiles and making informed healthcare decisions as they can reveal the complexity of in-vivo human phenotypic responses (Kuhn et al. 2010; Bowes et al. 2012). By understanding the underlying mechanisms of ADRs, we can gain insight into a drug’s mechanism of action, which can assist in identifying new drug targets, enhancing drug repurposing, predicting new therapeutic indications, and advancing personalized medicine.

Although there are ADRs that cannot be explained based on the known pharmacology or are due to the non-specific interactions of reactive metabolites, drug kinetics, and/or environmental exposures, most ADRs are caused by unintended consequences of on-target or off-target drug-protein interactions (Garon et al. 2017; Pirmohamed et al. 1998; Lounkine et al. 2012; Wallach, Jaitly, and Lilien 2010). Thus, drug target proteins serve as valuable resources for understanding the mechanisms of ADRs. Previous studies have considered the comprehensive set of drug targets to identify specific proteins associated with ADRs. An initial computational method for identifying ADR-related pathways (i.e., biological pathways that can explain the mechanisms of an ADR) was developed by Wallach et al. (Wallach, Jaitly, and Lilien 2010) who hypothesized that drugs modulating the same pathways may lead to ADRs with similar phenotypes. To establish ADR-pathway relationships, they employed a logistic regression model to predict ADRs by quantifying drug-pathway interactions based on the docking scores of drugs to proteins within each pathway. Mizutani et al. (2012) identified protein-associated ADRs by calculating the sparse canonical correlation between drug-protein relations and drug-side effect relations. Kuhn et al. (Kuhn et al. 2013) further defined the relationship of proteins to ADRs by searching for statistically significant overlap between the set of drugs linked to their associated proteins and the set of drugs linked to the given ADR. To establish the relationship between ADRs and their potential drug targets, Lounkine et al. (2012) calculated an enrichment score for each target-ADR pair based on their observed versus expected co-occurrence, and a statistical significance test was applied to find likely target-ADR associations. Lim et al. (2018) constructed a heterogeneous network including drug, gene, and ADR nodes. They employed the ADR-gene pairs identified by Lounkine et al. (2012) and applied a collaborative filtering-based algorithm to predict the missing links between ADRs and genes. Next, using a permutation-based algorithm, statistically significant genes for each ADR were ascertained and used for pathway enrichment analysis. Park et al. (2023) assumed that ADRs reported for drugs targeting a single protein are entirely derived from perturbing that specific protein. Accordingly, they hypothesized that predicting the likelihood that a single-target drug causes an ADR corresponds to the probability that the protein target is associated with the ADR. Based on this concept, they reduced the problem of ADR-protein association prediction to the problem of drug-single protein target prediction. To solve this problem, they constructed a network of drug-target and protein-protein interactions and used the node2vec representation algorithm to embed proteins and drugs into a low-dimensional vector space. They further used a logistic regression classifier for each ADR to score ADR-protein pairs.

Despite the importance of drug targets in understanding the mechanism of ADRs, exclusively relying on these protein targets can lead to false positives (or failure to identify the true causative pathway(s) owing to the limited search space). By contrast, exhaustive human genetic research has identified numerous genes related to diseases. These genes can often be linked to disease phenotypes that might be considered analogous to ADRs (Carss et al. 2022). Such relationships can be leveraged to strengthen our confidence and reduce the false positives complicating the drug target analysis approach. Nguyen et al (2019) hypothesized that phenotypes caused by genetic variations can be predictive of the phenotypes caused by drug interactions with the proteins encoded by those genes. They then showed that there is a statistically significant correlation between the organ systems affected by genetic variations and the organ systems exhibiting the ADRs when targeting the encoded protein(s).

In this study, we focus on the similarity between ADRs and disease phenotypes (DPs) by considering more specific phenotype MedDRA identifiers (low-level term), where MedDRA (Medical Dictionary for Regulatory Activities) is a medical terminology database used to standardize reporting of adverse drug reactions and other regulatory activities, and hypothesize that phenotypically similar ADRs and DPs might result from targeting of and variation in the same biological mechanisms and pathways (Fig. 1a). We expanded our analysis beyond individual genes/proteins by incorporating a protein-protein interaction network to consider the biological mechanisms. Relying on this hypothesis, we propose a novel network-based method to explore the mechanism of ADRs and DPs by identifying relevant proteins. Incorporating DP-associated proteins with ADR-associated proteins can reduce false positives compared to relying solely on those associated with ADRs. Moreover, by evaluating the same phenotypes in both ADRs and DPs, a more holistic view of the underlying molecular mechanisms can be unveiled.

**Fig. 1.**
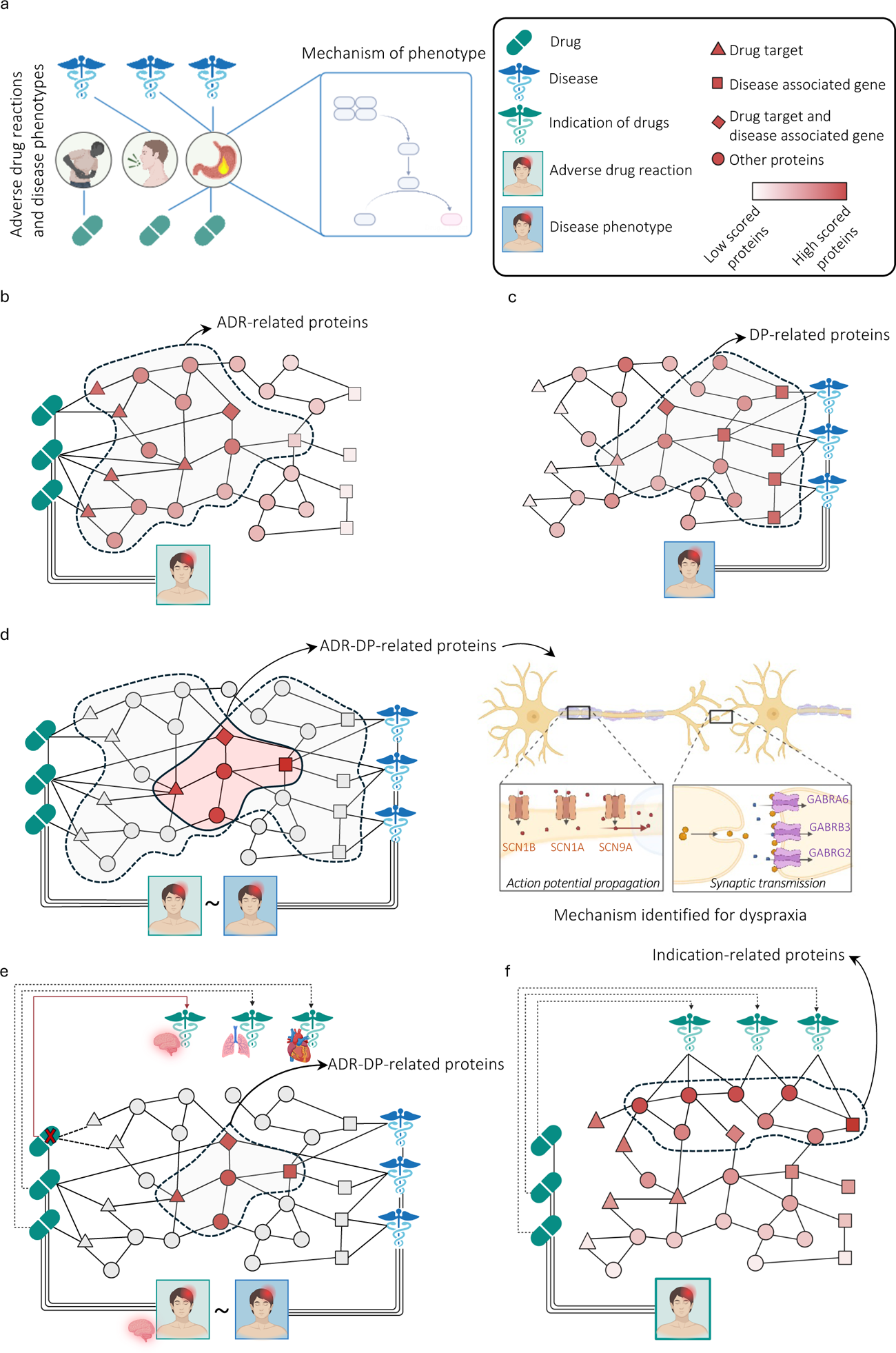
Overview of the DREAMER pipeline. (a) the basic hypothesis: phenotypically similar adverse drug reactions (ADRs) and disease phenotypes (DPs) might result from targeting of and variation in the same biological mechanisms and pathways; (b) to obtain ADR-related proteins, we diffuse from the drug targets and perform a statistical test for each protein; (c) similarly, to obtain Disease phenotype (DP)-related proteins, we diffuse from the disease related proteins and perform a statistical test for each protein. (d) left: ADR-DP proteins comprise the intersection set of proteins with significant overlap between ADR-related proteins and DP-related proteins; right: an example of identified ADR-DP proteins for dyspraxia (meddra.10009696) phenotype; three sodium channels (SCN1A, SCN9A, SCN1B) involved in action potential propagation and three GABA receptors (GABRB3, GABRG2, GABRA6) involved in synaptic transmission, all part of neuronal signaling. (e) to limit potential confounding effects by organ/tissue-related indications, ADR-DP proteins are identified after removing the drugs with the same organ/tissue indication as the organ/tissue affected by the ADR are excluded; and (f) to analyze further the confounding effects of drug indications, protein scores are determined by diffusing from proteins related to the indications of drugs associated with a specific ADR, resulting in the identification of significant proteins called indication-related proteins.

## 2. Results

### 2.1. Overview of the method

#### Network construction

We first combined several datasets to construct a heterogeneous network or multipartite network, also referred to as a knowledge graph (KG), in which drugs, diseases, proteins, adverse drug reactions (ADRs), and disease phenotypes (DPs) are represented as nodes. We first excluded ADRs linked to a large number of drugs (> 50), such as headache, nausea, skin rash, and diarrhea (Supplementary Fig. 1a), as well as DPs linked to a large number of diseases (> 100), such as seizure, hypotonia, and microcephaly (Supplementary Fig. 1b). We then linked pairs of ADRs and DPs based on the ADR-DP equivalence relationship deposited in the BioPortal database (Rubin et al., n.d.). Links between ADRs and drugs were obtained from the Sider database (Kuhn et al. 2013). We excluded links between ADRs and drugs that have indications equivalent to the ADRs in our datasets, which might be due to rare and unusual conditions or false positive reports. For example, metoprolol is a drug used to treat angina pectoris and hypertension while hypertension is listed as one of its ADRs. Links between DPs and diseases were collated from the Human Phenotype Ontology (HPO) database. We further linked drugs to proteins based on drug target information obtained from the DrugBank database (Wishart et al. 2018). In addition, diseases were linked to proteins encoded by disease-associated genes available in the DisGeNET database (Piñero et al. 2017). We also excluded drugs and diseases that had no known associations with proteins. Finally, the interactions between proteins were obtained from the STRING network (von Mering et al. 2003) with confidence scores above 0.8. While STRING encompasses a wide variety of interactions, including both direct (first order) and indirect (higher order) PPI interactions, we further analyzed the results using only physical interactions as the basis for constructing the protein-protein interaction network (Wang and Loscalzo 2021), which is more specific. Unless specified otherwise, the results presented in the main text are based on the STRING network. The framework used to prepare our KG is illustrated in Supplementary Fig. 2, and an overview of our KG is depicted in Supplementary Fig. 3. The summary statistics and data sources of our network are provided in Supplementary Tables 1 and 2, with more detail in the Methods section.

#### DREAMER pipeline

To investigate the underlying mechanisms of a specific ADR-DP pair, we first independently identified ADR-related proteins and DP-related proteins. To identify ADR-related proteins, we used a network diffusion algorithm, i.e., personalized page rank (see Methods) over the PPI network. As the initial condition, each protein in the network was assigned a probability based on the frequency of each protein being targeted by the drug associated with the queried ADR. Next, every protein score was allocated a p-value through a permutation test. Finally, proteins with corrected p-values less than 0.05 were considered ADR-related proteins (Fig 1b). To identify DP-related proteins, we followed the same approach replacing drug targets with disease-related proteins (Fig 1c). Finally, to minimize the false positives, we selected proteins at the intersection of the ADR and DP protein sets, so-called ADR-DP proteins – illustrated by the dyspraxia (meddra.10009696) phenotype as an example, involving six genes: three sodium channels (SCN1A, SCN9A, SCN1B) for action potential propagation, and three GABA_A_ receptor subunits (GABRB3, GABRG2, GABRA6) for synaptic transmission (Fig 1d, Supplementary Table 3). GABA_A_ receptors enhance sodium channel activation at myelinated axon nodes, regulating sensory feedback. Dysregulation can lead to dyspraxia due to impaired motor coordination. We excluded phenotypes for which no significant overlap between their ADR proteins and DP proteins was found (p-value < 0.05 – hypergeometric test adjusted by the Benjamini-Hochberg). The detailed description of our pipeline is explained in the Methods section. Moreover, to address potential confounding effects, we considered two aspects. First, indication-ADR organ overlap: removing drugs with the same organ/tissue indication as the ADR to avoid false associations (Fig. 1e). Second, indication-related proteins: scoring proteins by diffusing from those related to drug indications to identify and remove significant indication-related proteins (Fig. 1f, section 2.4).

As for visualization, we propose the diffusion map, which is a scatter plot representing each protein by its ADR-related and DP-related diffusion scores (Fig. 2a-c). The red points show proteins with scores that are statistically significant for both ADRs and DPs and usually have large diffusion scores for both ADRs and DPs. It is worth mentioning that proteins with high diffusion scores might not necessarily be significant. In certain cases, these proteins may be hub proteins (i.e., highly connected), enhancing the probability of achieving high scores in the null model and leading to their rejection in the permutation test. As predicted, this analysis identifies many proteins involved in disease processes. Next, we provide three examples of protein sets identified by DREAMER as significantly associated with ventricular arrhythmia, vasculitis, and thrombocytosis.

**Fig. 2.**
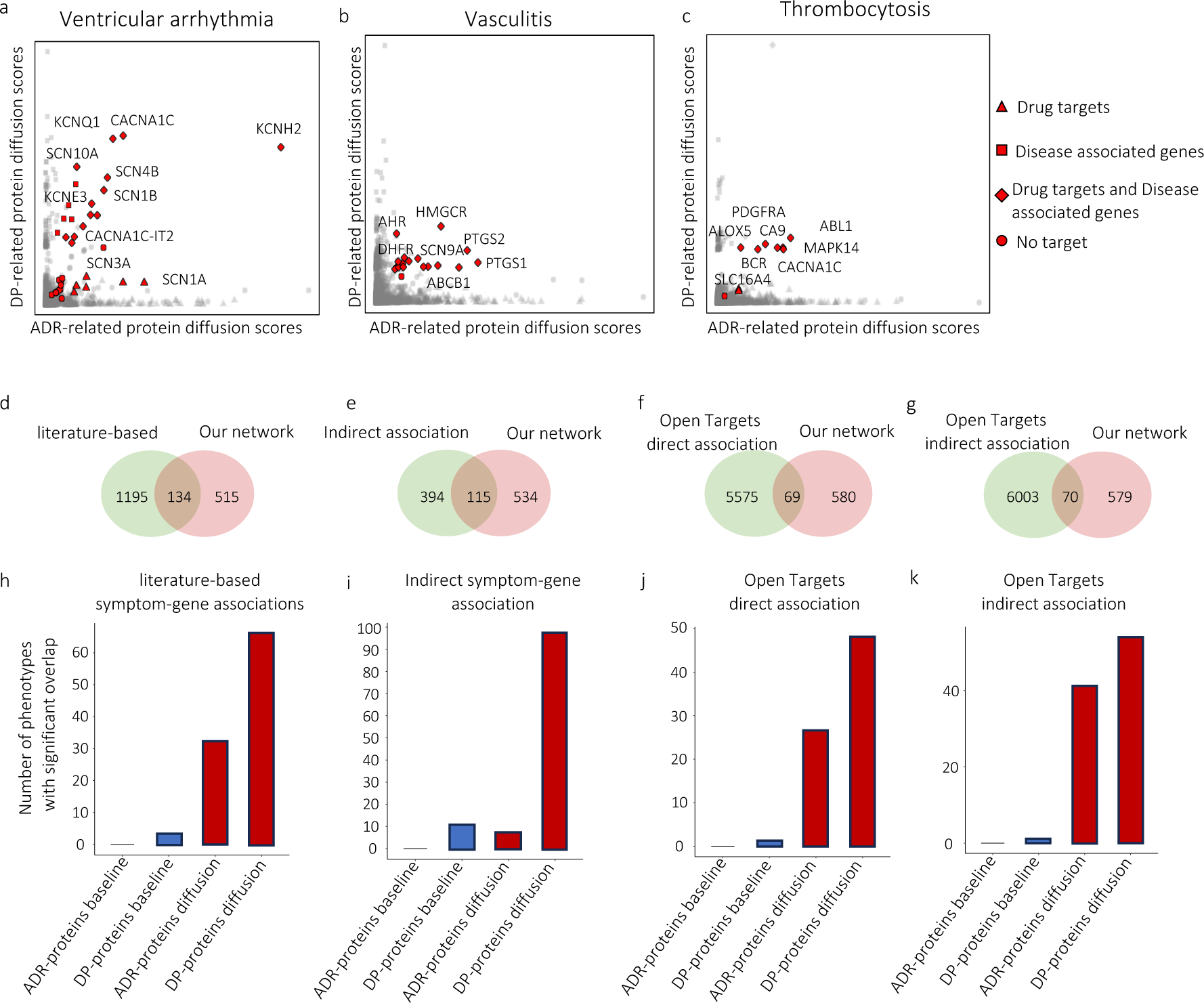
Diffusion map and reliability assessment of the identified protein set using the network diffusion algorithm on STRING; diffusion map for (a) ventricular arrhythmia, (b) vasculitis, and (c) thrombocytosis. The abscissa and ordinate values represent the diffusion scores of proteins initiated from the drug targets and disease-associated proteins, respectively. In the diffusion map, drug targets are represented by triangles, disease-associated proteins by squares, proteins that are both drug targets and disease-associated proteins by diamonds, and proteins that are neither drug targets nor disease-associated proteins by circles. Comparison of our diffusion-based method and the baseline method using (d, h) literature-based datasets, (e, i) indirect association based on disease-phenotype equivalent terms, (f, j) direct associations in Open Targets dataset, and (g, k) indirect associations in Open Targets dataset. d, e, f, g identifies the number of shared phenotypes with our constructed KG and known databases. h, i, j, k identifies the number of ADRs and DPs with significant overlaps between proteins identified by different methods and those reported in the known databases.

For ventricular arrhythmia (Fig. 2a), many of the significant proteins identified by the diffusion algorithm are ion channel proteins, such as those that contribute to Ca^2+^ (CACNA1C, CACNA1C-IT2), Na^+^ (SCN3A, SCN1B, SCN4B, SCN10A), or K^+^ (KCNQ1, KCNE3, KCNH2) transport in the heart (András et al. 2021). Previously described mutations in KCNQ1 and KCNH2 are associated with dysfunction of the voltage-gated K^+^ channel resulting in ventricular arrhythmias, such as long QT syndrome and ventricular fibrillation (Bellocq et al. 2004). Additionally, patients treated for ventricular arrhythmias often have their potassium (K^+^) levels tested and receive supplements if their levels are low. This is because hypokalemia, or low potassium levels, is a well-known risk factor for arrhythmias (Widimsky 2008). Mutations in Na^+^-channel proteins (SCN proteins) can result in long QT syndrome or atrial fibrillation (Platonov et al. 2019). Some of these ion channels are also present in other tissues, including brain, muscle, stomach, and colon. For example, mutations in the SCN1B can increase not only the risk of cardiac arrhythmia but also epilepsy (Baroni and Moran 2015). Therefore, drugs that alter their function can have cardiovascular, muscular, gastrointestinal, or neurological consequences, depending on which organs express the specific channels. Similarly, mutations in CACNA1C alter L-type voltage-gated Ca^2+^-channels and are associated with long QT and short QT syndromes. An example is Timothy syndrome, the complex congenital syndrome caused by CACNA1C mutations (Delinière et al. 2023), which involves cardiac manifestations such as long QT, along with one or more non-cardiac phenotypes such as skeletal, facial, and neurodevelopmental abnormalities (Borbás et al. 2022).

Vasculitis encompasses a heterogeneous group of diseases involving large, medium, or small vessels depending on the underlying specific disease (Acosta-Herrera et al. 2019; Pugh et al. 2022). Hallmarks include damage or dysfunction of the endothelial cells that line blood vessels, and treatments vary depending on the specific type. The disease phenotype reflects the action of specific proteins that govern the inflammatory response, including PTGS1 and PTGS2 (Fig. 2b), known as COX-1 and COX-2 enzymes. Kawasaki Disease, a pediatric vasculitis, is treated with aspirin targeting these enzymes and thereby reducing inflammation. Similarly, methotrexate, which inhibits DHFR (Fig. 2b), is used in the treatment of other forms of vasculitis, having more potent anti-inflammatory effects than aspirin or non-steroidal anti-inflammatory drugs (NSAIDs). Activation of AHR (Fig. 2b), the aryl hydrocarbon receptor, is also associated with promoting vascular inflammation; however, downregulation of AHR can also exacerbate vascular injury by enhancing the function of monocytes and macrophages (Wu et al. 2011; Nakajima et al. 2018). Statins, widely known for their ability to decrease cholesterol and reduce atherosclerosis via the inhibition of HMGCR (Fig. 2b), have beneficial anti-inflammatory effects on endothelial function and are being considered as additional therapies in some forms of vasculitis (Tremoulet 2015; Motoji et al. 2022).

Distinct from vasculitis, which involves inflammation of blood vessels, thrombocytosis is characterized by an elevated platelet count. This hematologic abnormality is reflected in the identified proteins that drive the phenotype, such as PDGFR-α and -β (platelet-derived growth factor receptor-alpha and beta) (Fig. 2c), which are present on both platelets and megakaryocytes, platelet precursors (Ye et al. 2010). Inhibition of these tyrosine kinases with imatinib and other related targeted therapies reduces megakaryocyte survival and proliferation and decreases platelet numbers by blocking PDGF signaling. Myeloproliferative syndromes, including essential thrombocythemia, can result from mutations in the JAK2, CALR, and MPL genes, each acting as drivers of the fusion protein BCR-ABL1 to increase cell (platelet as well as leukocyte) production (Kandarpa et al. 2024). The complete set of ADR-DP proteins for each phenotype is listed in Supplementary Table 3 and 4.

In summary, these examples demonstrate how the DREAMER pipeline can identify proteins that are mechanistically known to be associated with specific phenotypes.

### 2.2. Reliability assessment of the network diffusion method

In this section, we assess the reliability of network diffusion algorithms in identifying relevant proteins. To this end, we consider proteins that have previously been reported in the literature for each ADR or DP and calculate their overlap with ADR proteins and DP proteins, respectively. We further compare the ADR proteins and DP proteins with a baseline method that we call significant overlap association (see Methods section), previously used in the literature. Although there exists a limited number of curated resources of prior knowledge for ADR- and DP-related proteins (detailed below), they can still provide valuable sources to assess the reliability of our identified proteins and compare them to the baseline method. Leveraging these resources, we calculated the overlap between the computationally identified proteins (our method or the baseline method) and proteins obtained from *a priori* knowledge (known proteins). To determine the significance of these overlaps, we applied the hypergeometric test adjusted by the Benjamini-Hochberg test (p-value < 0.05). It is worth mentioning that both methods are applied to the same KG that we constructed.

#### Literature-based associations

Using the PubMed and SemMed literature databases, Lu et al. (2023) compiled DP-related proteins using natural language processing (NLP) methods and manual curation. They used NLP to search for the co-occurrence of DP and protein keywords through the text of abstracts in the published literature before January 2022. We found 134 phenotypes common to both the dataset of Lu et al. (2023) and our dataset (Fig. 2d, Supplementary Fig. 4a). We then assessed the significance of our network diffusion method over the baseline method in identifying proteins that have previously been found to be associated with each DP. To this end, we enumerate DPs whose related proteins have a significant overlap with those reported in the dataset (Fig. 2h, Supplementary Fig. 4b). As can be seen in Fig. 2h, being significant for 66 DPs, our identified proteins have a significantly larger overlap with those listed in the Lu et al. (2023) dataset compared to the baseline method. Interestingly, we also observe a relatively large number of overlapping proteins with our identified ADR-related proteins, although the Lu et al. (2023) dataset does not cover ADRs.

#### Indirect associations

In the datasets we used to construct our network, several DP terms are found to be equivalent to disease terms for which Genome-wide association studies-based related genes have been recognized. This equivalence can provide another opportunity for the assessment of our protein sets for such DP by defining their overlap with the set of proteins coded by the corresponding disease-related genes. To perform this analysis, we used the data provided by Lu et al. (2023) as they collected a set of DP-gene associations based on the similarity between disease and DP terms in the phenotype-genotype association databases. We found 115 DPs common to our dataset and the Lu dataset (Fig. 2e, Supplementary Fig. 4c). These results show the superiority of our method over the baseline method (Fig. 2i, Supplementary Fig. 4D).

#### Open Targets-derived data

The Open Targets platform has collected direct and indirect associations of targets and diseases from various sources, including genetic associations, somatic mutations, drugs, pathways, RNA expression, text mining, and animal models (Carvalho-Silva et al. 2019). Direct association focuses on the evidence that specifically refers to the target and the phenotype in question, such as whether multiple studies directly link the NOD2 gene to inflammatory bowel disease. Indirect association, by contrast, leverages the hierarchical structure of the disease ontology, allowing the inclusion of relevant evidence that may not explicitly mention the primary disease or phenotype of interest but may still be pertinent owing to the related nature of the conditions (for example, when assessing the association between inflammatory bowel disease and NOD2, evidence linking *Crohn disease*, a descendant in the disease ontology of inflammatory bowel disease, to NOD2 is also considered). Applying this dataset, we found 69 phenotypes with direct associations and 70 with indirect associations common to Open Targets and our dataset (Fig. 2f and 2g, Supplementary Fig. 4e and 2g), which we considered for the assessment of our pipeline (Fig. 2j and 2k, Supplementary Fig. 4f and 2h). As can be seen, our diffusion-based method outperforms the baseline method in identifying proteins reported in the Open Targets dataset. It is worth noting that none of the disease-gene associations in the Open Targets dataset was found in our KG.

In our validation analysis, we note that we do not expect a perfect overlap between our recognized protein sets and literature-based proteins. This lack of complete consonance is a consequence of the fact that the available literature-based protein sets are not sufficiently comprehensive. Therefore, although showing some degree of overlap of ADR-related and DP-related proteins with proteins derived from literature- and genetic-based sources is useful for validation, we expect to recognize proteins *de novo* for each phenotype. Moreover, to reduce the false positives from ADR proteins and DP proteins, we identified the intersecting proteins between these two groups. Among 649 phenotypes, 120 of them had a significant number of overlapping proteins between their identified ADR proteins and DP proteins (p-value < 0.05 – hypergeometric test adjusted by the Benjamini-Hochberg test), an overlapping protein set that we term ADR-DP proteins. Hence, we consider these 120 protein sets for the downstream tasks.

### 2.3. Holdout validation

DREAMER identifies a set of proteins mechanistically related to a phenotype by analyzing the network proximity of proteins with drugs and diseases that are associated with that phenotype. This outcome can raise the question of generalizability of the proteins identified by DREAMER when exposed to new drugs or diseases that were not included during the discovery phase, i.e., the process of identifying proteins by DREAMER. Accordingly, we hypothesized that a new drug or disease associated with a phenotype that was not included in the discovery phase should probably have at least one associated protein closer to the identified proteins than drugs and diseases that do not have that phenotype (Supplementary Fig. 5). To test this hypothesis, we performed a holdout analysis to validate further our proposed pipeline. In particular, we divided the drugs and diseases associated with each phenotype into two groups: 80% for discovery and 20% for validation. For each phenotype, we also randomly selected drugs and diseases that are not associated with the given phenotype, with an equal number as in the validation set. Here, the drugs or diseases in the discovery set are referred to as positive assets, and those randomly selected are so-called negative assets. In the discovery phase, DREAMER identified proteins related to each phenotype using the discovery set. We then obtained the shortest paths from the identified proteins to the positive sample assets, as well as the negative assets.

For positive and negative drugs (and diseases), we counted the number of shortest paths with lengths equal to or less than X ∈ {0, 1, 2, …} across all phenotypes. Using Fisher’s exact test, we investigated if the ratio of positive assets with the shortest path ≤ X is significantly larger than those of negative assets. The results of the holdout analysis of drugs and diseases are shown in Table 1. As can be seen, the drugs (or diseases) in the validation set are significantly closer to the proteins identified by DREAMER for each phenotype compared to randomly selected drugs (or diseases), which supports our hypothesis that phenotypically similar ADRs and DPs might result from targeting of and variation in the same proteins and pathways.

**Table 1.**
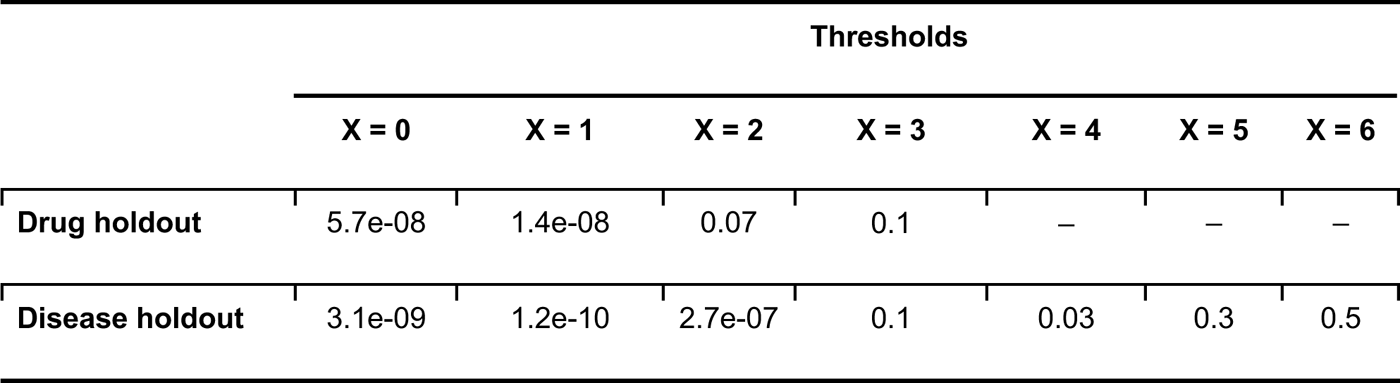
The p-values obtained from Fisher’s exact test for the holdout validation.

Additionally, we repeated the process above by splitting the drugs into discovery and validation sets based on their molecular similarities using the *DataSAIL* package in Python (Joeres, Blumenthal, and Kalinina 2023). In particular, to split the molecules into discovery and validation sets, *DataSAIL* uses an algorithm to minimize the similarity between the molecules in the discovery set and those in the validation set. To obtain the similarity between drugs, *DataSAIL* uses Tanimoto coefficients between molecular fingerprints calculated using their Simplified Molecular Input Line Entry System (SMILES) designations. This approach ensures that the drugs in the discovery and validation sets are not structurally similar, thereby preventing information leakage. The results are presented in Supplementary Table 5, which are in line with the results presented above.

### 2.4. Considering the confounding effect of drug indications

The proteins identified by DREAMER for a specific phenotype may be influenced by the effects of indications of drugs associated with the corresponding ADR. In this context, we discuss two types of potential confounding effects:

[i] drugs whose indication affects the same organ/tissue as their ADR might have targets that seem relevant to the ADR; however, these associations could be only due to the drug’s primary indication rather than to the underlying mechanism of the ADR; and
[ii] indications of drugs associated with the corresponding ADR might be associated with proteins that are closely related to the ADR-DP protein set (as identified by DREAMER); in this case, it is possible that the observed relationship with ADR-proteins is driven by the indications rather than by a direct connection to the ADR.

In the following paragraphs, we focus on each of these confounding effects and describe the pipeline employed to address them.

To limit potential confounding by organ/tissue-related indications in our analysis, we excluded drugs with the same organ/tissue indication as the organ/tissue affected by the ADR (Fig. 1e). To this end, we manually identified the involved organs/tissues for the 120 ADR-DPs, discussed in section 2.2 (Supplementary Table 6), and we obtained organ indications of drugs from Nguyen et al. (2019). Among 465 drugs in our network, 328 drugs were listed in their dataset. Drugs that have indications and associated ADR of the same organ/tissue were subsequently excluded. For example, for the cardiovascular ADR phenotype, tachycardia, we eliminated all of the associated drugs with cardiovascular indications. Subsequently, the number of ADRs with at least one associated drug was reduced to 97 (listed in Supplementary Tables 7 and 8). Next, we applied the DREAMER pipeline to obtain a new set of proteins for each phenotype. For 90 out of 97 phenotypes, we observed a significant overlap between the proteins identified before and after the organ/tissue-based drug removal described above (tested via hypergeometric test, p-value < 0.05) listed in Supplementary Tables 7 and 8. We note that, after the removal of the drugs with indications in the same organ/tissue as the ADRs, the average number of drugs was reduced to 6.6 from 14.2 for each ADR. While, on average, 30% of the drugs are excluded in this analysis, the results do not show a significant variation with the analysis in which this filter was not applied, confirming the robustness of our pipeline.

We further sought evidence that the ADR-DP proteins for each phenotype interact with proteins associated with indications (so-called indication-related proteins) of drugs related to the corresponding phenotype. To this end, we used the network diffusion algorithm over the PPI network (as we used for identifying ADR and DP proteins) to assign a diffusion score to proteins with respect to their PPI adjacencies with the indications of drugs (Fig. 1f). The diffusion algorithm was initialized based on the frequency of proteins associated with these indications. Among 120 phenotypes (see section 2.2), 95 of them were linked to at least one drug with at least one indication that has at least one associated gene listed in our dataset (listed in Supplementary Tables 7 and 8). Restricting our analysis to these 95 ADRs, we obtained diffusion scores from the indications of drugs linked to each ADR. Fig. 3 shows the diffusion map now incorporating a third dimension representing the diffusion scores by drug indications. Next, every protein score was allocated a p-value through a permutation test, and proteins with corrected p-values less than 0.05 were considered as indication-related proteins (see Methods section 4.2 and 4.3). We then recognized phenotypes with significant overlap (hypergeometric test with Benjamini-Hochberg adjusted p-value < 0.05) between the indication-related proteins and ADR-DP proteins. For 84 out of 95 phenotypes (listed in Supplementary Tables 7 and 8), no evidence of significant overlap was found. As can be seen, in these phenotypes, the ADR-DP proteins (indicated in red) have small diffusion scores with respect to the third dimension, suggesting that for these phenotypes, the subnetworks related to drug indications are far from those related to ADR-DP proteins. Examples of these phenotypes are shown in Fig. 3a–c. For example, intracranial hemorrhage (Fig. 3c), a critical condition involving bleeding within the brain, was linked to proteins of the coagulation and anticoagulation pathways such as PROC (protein C), F10 (factor X), and F2 (prothrombin) (Supplementary Table 3). Dysregulation of these proteins can impair hemostasis and form stable clots, leading to an increased risk of excessive bleeding events such as intracranial hemorrhage. These findings suggest that, at least for these phenotypes, the identified ADR-DP proteins have no significant association with the protein drivers of their clinical indications for their related drugs.

**Fig. 3.**
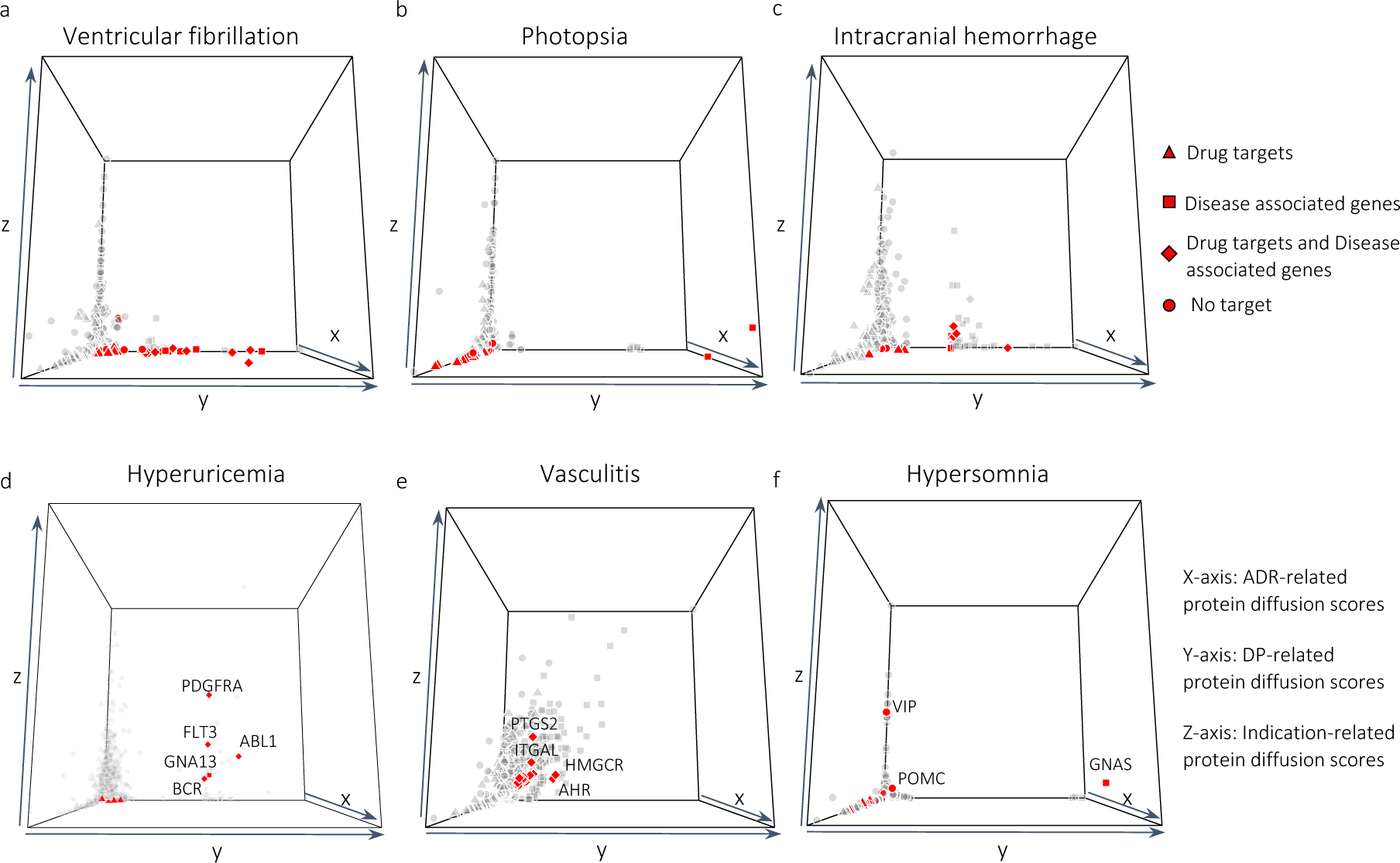
3D diffusion map; x, y, and z-axis represent the diffusion scores of proteins from drug-targets, disease-protein, and drug indication-protein, respectively for (a) ventricular fibrillation, (b) photopsia, (c) intracranial hemorrhage, (d) hyperuricemia, (e) vasculitis, and (f) hypersomnia. The ADR-DP proteins are indicated as red points.

By contrast, for the remaining 11 phenotypes (Fig. 3d–f and Supplementary Fig. 6), the ADR-DP proteins have larger values in the third axis. For these phenotypes, the independence of ADR-DP proteins and the indication-related proteins is not trivial and an additional step of investigation by domain-specific experts is required. Accordingly, the 3D diffusion map can be used to inspect any ambiguity among the identified proteins. Put another way, ADR-DP proteins with a high value on the z-axis may reflect reverse causality—i.e., the ADR is a downstream consequence of treatment by indication and may not “directly” be associated with the underlying mechanism of the associated phenotype. For example, in hyperuricemia (Fig. 3d), we identified ADR-DP proteins that included PDGFRA, FLT3, and ABL1 as having high values in the z-axis. These proteins are commonly targeted by drugs for cancer treatment, including leukemia, and play roles in the differentiation, division, and growth of cells. Cell death induced by these treatments can lead to great increases in uric acid in the blood, overwhelming the body’s normal ability to clear that metabolite and ultimately causing renal dysfunction further worsening the hyperuricemia. For vasculitis, as discussed in section 2.1, we have identified proteins, such as PTGS2 (Fig. 2b), that are crucial in mediating inflammation and pain via prostaglandin synthesis. Incorporating indication-related proteins, we observe an overlap in inflammatory activation between the indication (z-axis) and vasculitis (y-axis) (Fig. 3e). Similarly, ITGAL (CD11a/LFA-1) (Fig. 3e), essential in leukocyte migration (Ley et al. 2007), may be detected due to the (i) modulation of immune filtration by the indicated drug/disease and/or (ii) vasculitis itself affecting immune cell-endothelial interactions. For hypersomnia, VIP’s high value in the z-axis might reflect its connection to VIPomas, where octreotide inhibits excessive VIP secretion (Fig. 3f). However, VIP is produced by neurons in the suprachiasmatic nucleus (SCN) of the hypothalamus where it maintains normal circadian rhythms (Todd et al. 2020), supporting its potential involvement in hypersomnia too. VIP’s role in various phenotypes likely depends on its anatomic location and quantification.

According to both analyses discussed above, we identified mechanisms for 67 phenotypes that show no evidence of association with drug indications (Supplementary Table 7). Additionally, when replacing the STRING PPI network with the physical PPI network, our analysis identified mechanisms for 56 phenotypes (Supplementary Table 8) that are not related to drug indications. Notably, there was an overlap of 29 phenotypes between these two analyses. The ADR-DP proteins identified from both analyses for all these 29 phenotypes showed significant overlap (hypergeometric test, adjusted using the Benjamini-Hochberg method, p-value < 0.05).

### 2.5. Biological insights into the phenotype mechanism of action

In this section, we investigate the biological function of the protein sets identified by DREAMER for 67 phenotypes with no evidence of association to drug indications, as discussed in section 2.4. We first ranked these phenotypes based on the significance of the overlaps between their ADR proteins and DP proteins, which can be viewed as an indicator of their alignment with our initial hypothesis. Supplementary Fig. 7 shows the ranking of all phenotypes, with the top 20 phenotypes shown in Fig. 4a. We then found the enriched pathways for the ADR-DP protein sets based on an over-representation analysis using the Reactome (Croft et al. 2011) and Gene Ontology (GO –(Harris et al. 2004)) databases. The results for the top three ranked phenotypes are illustrated in Fig. 4b–g, and a complete set of the enriched pathways for each phenotype is listed in Supplementary Table 9.

**Fig. 4.**
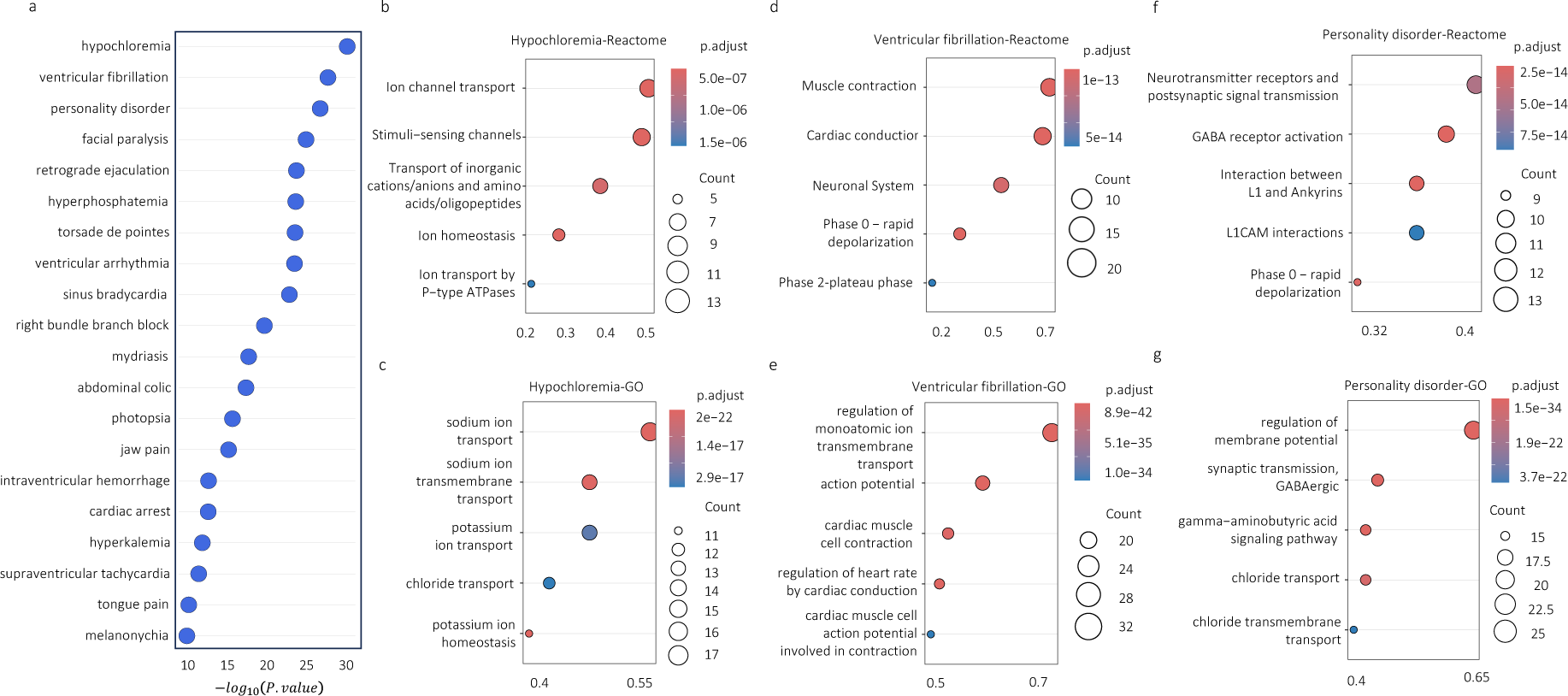
The results for the top phenotypes; (a) The ranked list of phenotypes based on the significance of their ADR-DP proteins. Reactome (b, d, f) and gene ontology (c, e, g) over-representation analysis for ADR-DP proteins of the top three phenotypes including (b, c) hyperchloremia, (d–e) ventricular fibrillation, and (f–g) personality disorder.

Pathway analysis highlights the physiological processes involved in these disorders. For example, chloride is an anion that is mostly found in the extracellular space. Its concentration is regulated by the gastrointestinal tract where it is absorbed from food, as well as the kidney where it is excreted in urine or reabsorbed in the proximal tubule. Chloride transport relies on transmembrane ion transporters and cotransporters as well as additional Na^+^/K^+^ ATP-dependent ions transporters that provide energetics for Cl^-^ transport (Fig. 4b-c). Thus, its concentration is dependent on that of other ions, such as Na^+^, K^+^, and bicarbonate (HCO_3_^-^). Owing to its inverse relationship with bicarbonate, hypochloremia can result in metabolic alkalosis. Hypochloremia can occur due to gastrointestinal causes, such as vomiting; or renal loss of chloride due to the use of diuretics (hypochloremic metabolic alkalosis due to excessive fluid loss leading to volume contraction) and/or because of hyponatremia and hypokalemia, as the fluxes in sodium and potassium will affect chloride levels (Berend, van Hulsteijn, and Gans 2012).

The coordinated movement of ions through voltage-gated ion channels is important to maintain the rhythmic beating of the heart (Fig. 4d-e). Disruption of these processes leads to abnormal action potentials, arrhythmias, and ventricular fibrillation. Many of these ion channels also play a role in other organs, including the brain (Niemeyer et al. 2001).

In personality disorder (a complex, comparatively nonspecific phenotype), the identified pathways are all known key mechanisms for various psychiatric conditions. The neurotransmitter receptors and postsynaptic signal transmission reflect the significant roles of dysregulated neurotransmitter systems implicated in a range of personality disorders such as mood and bipolar disorder. Altered phase 0, representing rapid neuronal depolarization, can lead to epilepsy (Holmes and Ben-Ari 2001) (Fig. 4f). Dysregulated membrane potential can be influenced by chloride transport and can impair GABAergic transmission (Fig. 4g). Impairment in GABAergic transmission plays a significant role in the pathophysiology of major depressive disorder (MDD) (Luscher, Shen, and Sahir 2011), schizophrenia (Gonzalez-Burgos, Fish, and Lewis 2011), bipolar disorder (Brambilla et al. 2003), and autism (Horder et al. 2018), and lower levels of GABA are often identified as the main endophenotype of MDD (Hasler and Northoff 2011). The antidepressant effect of ketamine may also be related to its selective impact on GABAergic interneurons, blocking NMDA receptors and reducing inhibitory signals to enhance cortical excitation. Additionally, the interaction between L1CAM and ankyrins (Fig. 4f) guides neuronal adhesion and signaling, where abnormalities are associated with neurodevelopmental disorders like autism (Yang et al. 2019). These mechanistic phenotype pathways emphasize the interconnected roles of neurotransmitter signaling, synaptic function, and neuronal excitability in personality (and other psychiatric) disorders.

### 2.6. Application of DREAMER in exploring therapeutic potential

The proteins identified for each phenotype using DREAMER can open new avenues for drug design and drug repurposing, i.e., an approach to identifying new therapeutic uses for drugs that are already approved for specific disorders (Pushpakom et al. 2019). It can be hypothesized that targeting proteins identified for each phenotype is most likely either to induce or treat the phenotype, as one cannot determine directionality *a priori* from this analysis. Therefore, DREAMER can be leveraged for drug discovery in two ways: [i] predict possible ADRs for new drugs based on their known targets, and [ii] design new drugs or suggest repurposing candidates, based on their targets, for a disease.

In particular, to showcase the application of DREAMER for drug repurposing in the context of the second case, we focus on phenotypes for which there is evidence that targeting their ADR-DP proteins can treat the corresponding phenotype. For this purpose, we identified phenotypes whose ADR-DP proteins contain at least one protein targeted by a drug with an indication with the same terminology as the specified ADR. For example, cardiac arrest is a terminology that is assigned to an ADR (with meddra:10007515 in the SIDER dataset), a disease phenotype (with hpo:0001695 in the HPO dataset), and drug indication (with mondo:0000745 in Mondo Disease Ontology dataset). Interestingly, we found three drugs, diltiazem, carvedilol, and verapamil, that are indicated for cardiac arrest and target at least one of the proteins recognized by DREAMER for cardiac arrest (Fig. 5a). Thus, hereafter we refer to such drugs as “indicated drugs”. In our dataset, we identified a total of eight such phenotypes including cardiac arrest, hypophosphatemia, precocious puberty, torsade de pointes, thrombocytosis, peptic ulcer, ventricular tachycardia, and ventricular fibrillation (Supplementary Table 10). Fig. 5 illustrates PPI subnetworks for five of those phenotypes restricted to their ADR-DP proteins along with the drugs that target them and have indications for those phenotypes. The complete list of the indicated drugs along with their targets among ADR-DP proteins are provided in Supplementary Table 10.

**Fig. 5.**
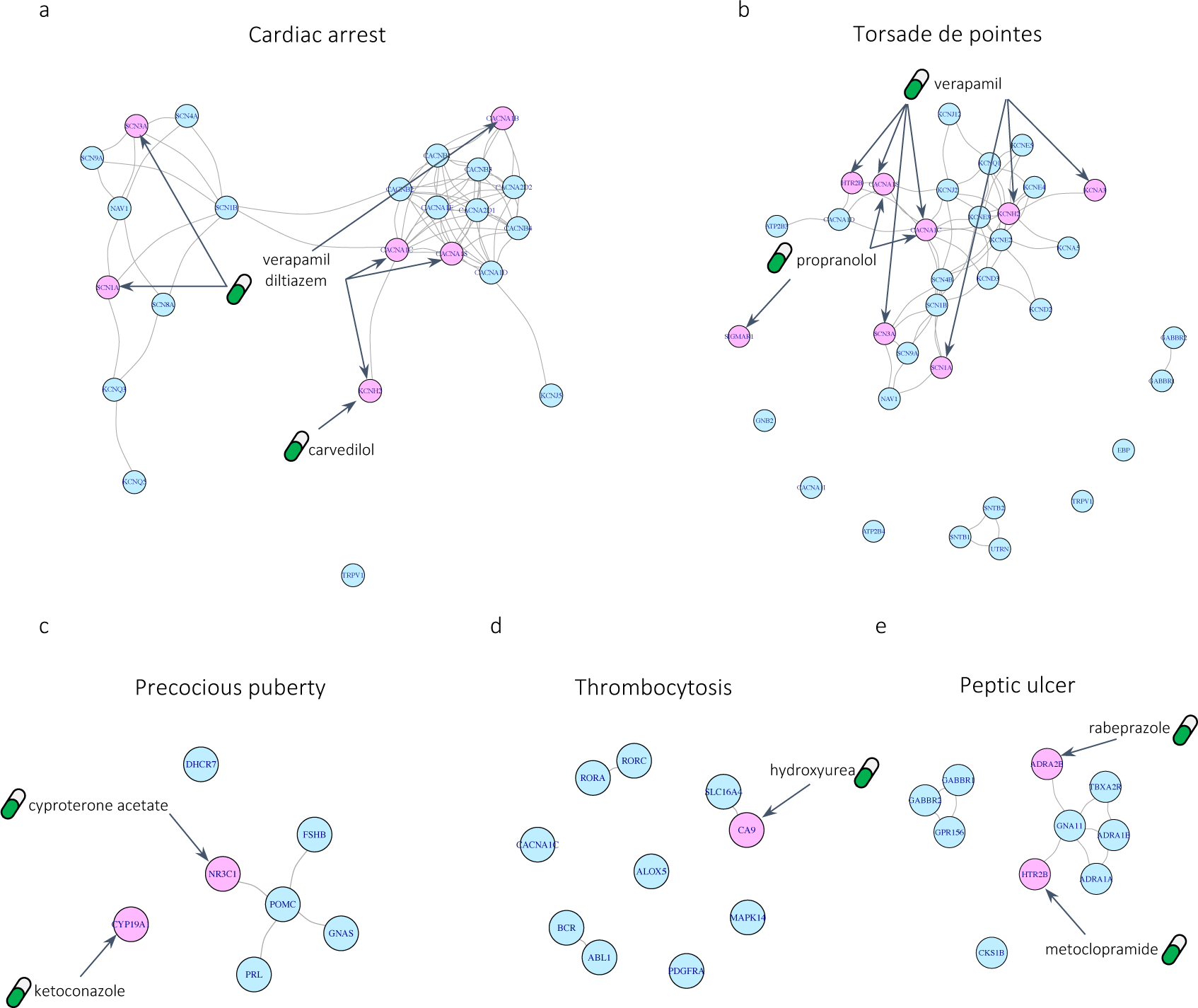
Subnetworks of identified protein sets along with indicated drugs that target them; (a) precocious puberty, (b) thrombocytosis, (c) peptic ulcer (d) torsade de pointes, and (e) cardiac arrest. Pink nodes are the proteins that are targeted by indicated drugs and blue nodes are the rest of the ADR-DP proteins.

To find opportunities for drug repurposing on the above mentioned phenotypes, we found all the drugs that have at least one target among their ADR-DP proteins and focused only on those with no ADR on the corresponding phenotype and those that have not previously been found to have an indication for the corresponding phenotype in our dataset (Supplementary Table 11). We, thus, refer to these drugs as “candidate drugs” for repurposing. For example, sotalol is recognized for its efficacy in treating various cardiac arrhythmias by targeting KCNH2 (hERG) channels. In our dataset, sotalol is recognized to have indications for ventricular fibrillation. However, sotalol can cause prolongation of the QT interval, leading to ventricular arrhythmias such as ventricular tachycardia, ventricular fibrillation, cardiac arrest, and, in particular (based on our dataset confirmed by the published literature), torsades de pointes (Kpaeyeh and Wharton 2016; Wharton et al. 2022; Hohnloser and Woosley 1994). Although sotalol has not been reported to be related to cardiac arrest in our dataset, it is among the candidate drugs for the treatment of cardiac arrest in our analysis (Supplementary Table 11). Similarly, ranolazine, a drug used to treat angina pectoris, has been used off-label for the treatment of ventricular arrhythmias (Andrade and Deyell 2022). In addition, we conducted a comprehensive search on the clinical trials website (*clinicaltrials.gov*) to find evidence for these candidate drugs. Using a customized Python script, we queried all pairs of phenotypes and their candidate drugs, and then carefully inspected all of the derived results. Accordingly, we found a number of such phenotype-drug pairs listed in Table 2 along with their ClinicalTrials.gov IDs.

**Table 2.**
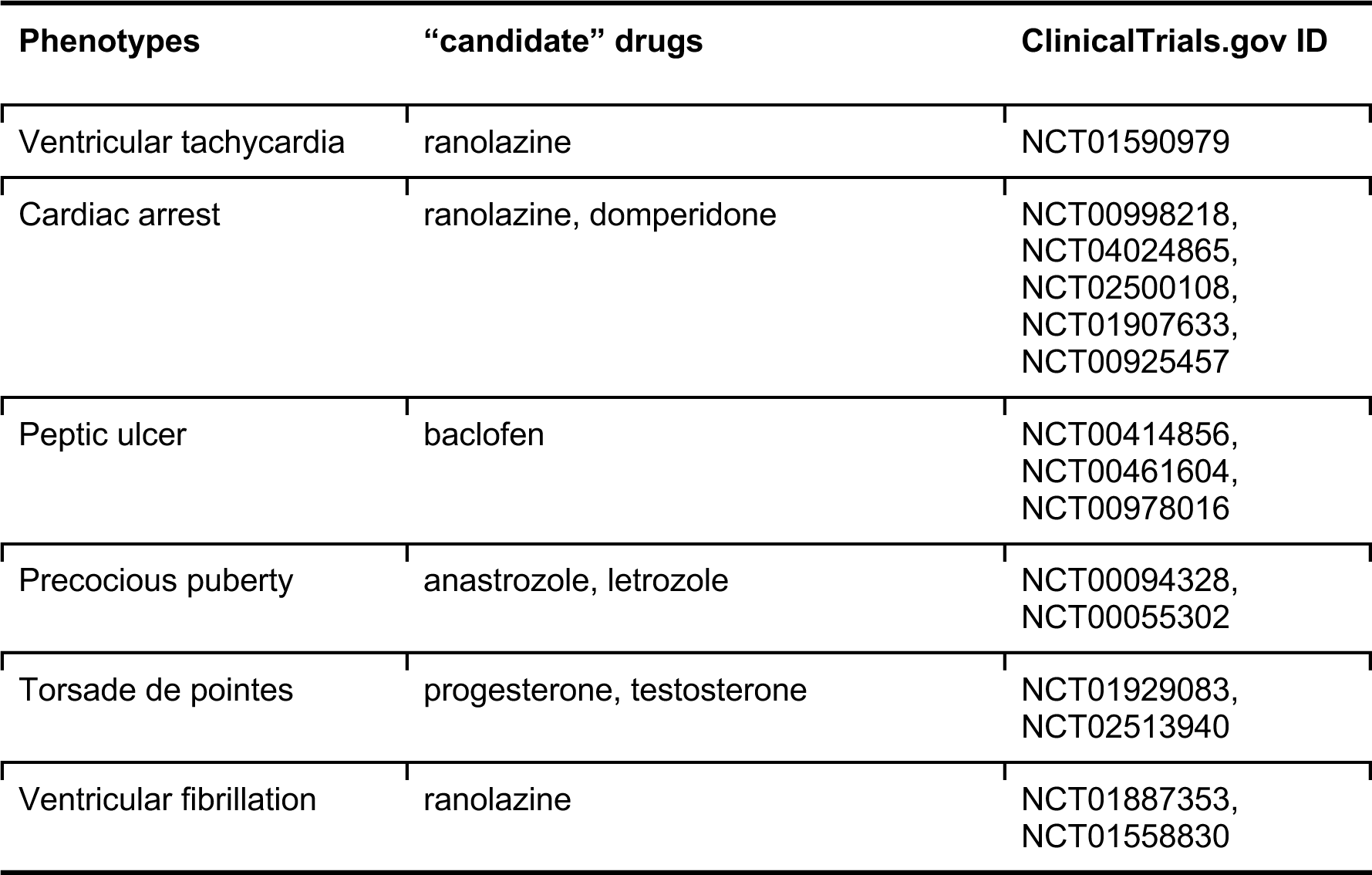
The phenotype-drug evidence for “candidate” drugs in the clinical trials website.

| Phenotypes | “candidate” drugs | ClinicalTrials.gov ID |
| --- | --- | --- |
| Ventricular tachycardia | ranolazine | NCT01590979 |
| Cardiac arrest | ranolazine, domperidone | NCT00998218,<br>NCT04024865,<br>NCT02500108,<br>NCT01907633,<br>NCT00925457 |
| Peptic ulcer | baclofen | NCT00414856,<br>NCT00461604,<br>NCT00978016 |
| Precocious puberty | anastrozole, letrozole | NCT00094328,<br>NCT00055302 |
| Torsade de pointes | progesterone, testosterone | NCT01929083,<br>NCT02513940 |
| Ventricular fibrillation | ranolazine | NCT01887353,<br>NCT01558830 |

To repurpose novel drugs for these phenotypes, we scored and ranked all candidate drugs based on the ratio between the number of proteins they target within ADR-DP proteins to their overall number of targets. Fig. 6 shows these rankings for the top 20 drugs, with circle sizes representing the total number of targets of each drug within the ADR-DP protein set. The circles are colored “red” to indicate drugs that have been found to be under investigation in clinical trials as being relevant to the queried phenotype. Candidate drugs that target proteins also targeted by indicated drugs may have a potential for repurposing (marked in “green”), as targeting these proteins has already been shown to be effective in treating the phenotype. Circles colored in “blue” could potentially represent interesting findings to repurpose for their corresponding phenotypes, as they have not been previously targeted for the treatment of those phenotypes. Moreover, we hypothesize that drugs with higher ranks are more likely associated with the corresponding phenotype. Nonetheless, experimental and clinical trials are essential for their validation.

**Fig. 6.**
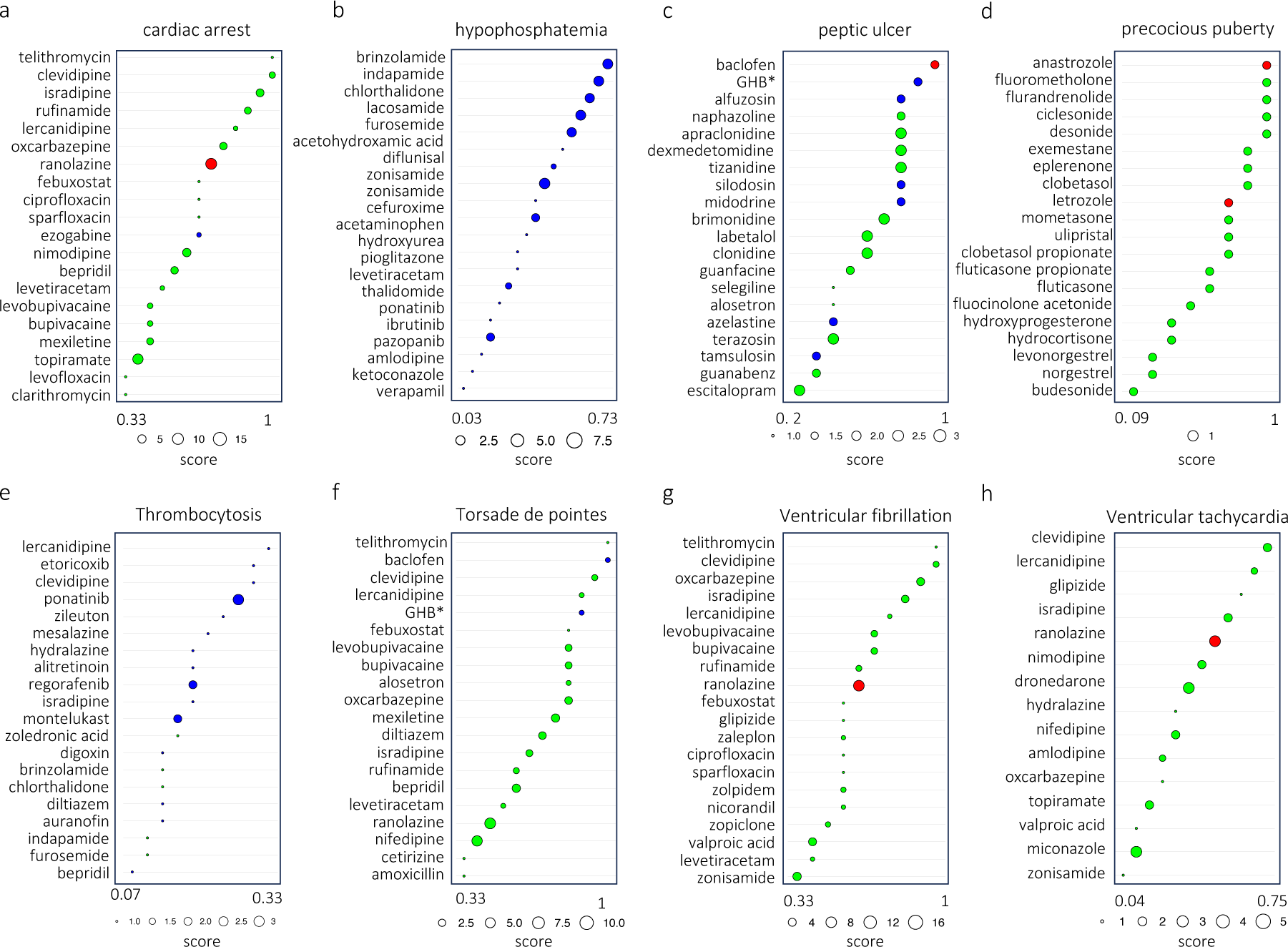
Top-ranked drugs with at least one target in the identified ADR-DP proteins for the phenotypes; (a) cardiac arrest, (b) hypophosphatemia, (c) peptic ulcer, (d) precocious puberty, (e) thrombocytosis, (f) torsade de pointes, (g) Ventricular fibrillation, and (h) ventricular tachycardia. Abbreviation: GHB* for gamma−hydroxybutyric acid. “Red” circles indicate candidate drugs that are found to be under investigation in clinical trials. “Green” circles indicate candidate drugs that target proteins that are already targeted by indicated drugs. “Blue” circles indicate candidate drugs whose targets were not already targeted by any indicated drugs. The size of the circles indicates the total number of proteins the candidate drugs target within ADR-DP proteins.

## 3. Discussion

In this study, we present DREAMER, a network-based method to investigate the underlying mechanisms of DPs and ADRs. Although there have been previous efforts to investigate these mechanisms separately, focusing on ADR-target protein interactions or DP-gene associations, utilizing both phenotypes together offers a more comprehensive insight into their mechanisms. By elucidating the interconnected modules between ADRs and DPs, our approach has the potential to explore biomarkers for designing safer therapeutic strategies that minimize the need for drug discontinuation and enhance the potential for drug repurposing, thereby ensuring more effective and personalized treatment modalities.

Targeting proteins in the organ of interest with drugs provides the basis for *in vivo* experiments that can explain the relationship between the functionality of that protein, systemic effects, and phenotypic responses. These proteins can be both on-target and off-target of drugs. While they are primarily targeted to treat specific indications or their phenotypes, they can also induce unintended side effects. Currently, drug safety evaluation, to a large degree, relies on animal experiments, which may not always translate reliably to humans owing to inherent biological differences (Carss et al. 2022). In recent years, the increased availability of public databases including drug targets and ADRs has become a more reliable source of human-specific information. Genetic variations, by contrast, can be considered natural experiments, providing insights into the mechanism of phenotypes. Genome-wide association studies have been extensively utilized to identify novel therapeutic targets, with a greater probability of drug approval when these targets are corroborated by human genetic evidence for the desired indication. Additionally, there is a growing interest in harnessing human genetic studies to predict the risk of ADRs. The importance of applying this strategy is more pronounced where suitable animal models for drug safety assessment are lacking (Carss et al. 2022). Protein modules that are affected by both drugs with a specific ADR and diseases with a similar phenotype provide more evidence to explain the mechanism underlying the phenotype. Our study advances this concept by considering modules targeted by drugs and diseases exhibiting phenotypically similar ADRs and DPs.

The ADR-DP proteins identified by DREAMER may be influenced by the drug indications in the PPI, which can be recognized by their large values along the z-axis in the 3D diffusion map (Fig. 3). Such instances may affect our interpretations and should be addressed carefully. Specifically, we encountered three scenarios: [i] ADR-DP proteins have an indirect association with the phenotype of interest, as seen in cases like hyperuricemia caused by cancer therapies (refer to section 2.4); [ii] higher-order relationships between the drug indications and the phenotype (such as both being related to the same tissue or organ); and [iii] the phenotype mechanism stems from on-targets effects. We acknowledge that while the last scenario does not limit our interpretation, distinguishing it from the first two scenarios might be challenging or even infeasible.

Some phenotypes are multifactorial and are not directly linked to the drug’s molecular effect(s) alone. However, it is still valuable to explore whether any of these phenotypes have a specific molecular mechanism that connects them to both ADRs and DPs simultaneously. For example, in the case of female infertility, which often stems from prior infections, DREAMER has identified proteins enriched in the metabolism of steroid hormone pathways (Supplementary Table 3 and 9). In contrast, dementia may arise from multiple factors beyond genetics, such as age, lifestyle, social engagement, and cognitive function, which would be discarded by DREAMER owing to the lack of significant overlapping proteins connecting the ADRs and DPs.

In this analysis, we used the STRING network, which encompasses a wide variety of protein-protein interactions from various sources. This integrative network allows for the exploration of biological interactions, including direct physical interaction as well as indirect (higher-order) associations such as genetic co-occurrences, co-expression, and others derived from computational predictions and existing literature. Such a diverse compilation of interaction types is useful to elucidate the phenotype mechanisms based on higher-order and wider ranging interactions. Despite the advantages offered by the STRING database in uncovering potential phenotype mechanisms through its generalized interaction data, we also found it helpful to apply DREAMER further on the basis of a physical interaction network alone, which includes interactions validated by high-confidence experimental methods such as yeast two-hybrid screening, NMR spectroscopy, x-ray crystallography, and cryo-electron microscopy. The shared phenotypes (physical PPI network and STRING) are extensively studied, commonly encountered in clinical practice, and cover a broad spectrum of organ systems, such as cardiac arrest, bradykinesia, ventricular arrhythmia, and interstitial pneumonitis (Supplementary Fig. 7). For neurological and muscular abnormalities, both the physical PPI network and STRING datasets detect bradykinesia, facial paralysis, hemiplegia, muscle stiffness, and mydriasis. The physical PPI network, however, uniquely captures dementia (cognitive symptoms) and parkinsonism (cognitive and motor symptoms), whereas STRING uniquely identifies akinesia, a symptom of parkinsonism related to movement initiation difficulties. The reasons for these distinctions remain uncertain at this time, but likely reflect the complexity of the phenotypes as well as their lack of specificity in some cases (e.g., dementia). This trend is also observed in inflammatory phenotypes. Both methods identify interstitial pneumonitis, an inflammation of lung tissue. However, the physical PPI network uniquely identifies systemic inflammatory disease phenotypes, such as rheumatoid arthritis and leukocytosis, involving direct interactions with mediators derived from circulating immune cells, while STRING identifies organ-related inflammation through broader network interactions, such as cholecystitis and cholangitis (Supplementary Fig. 7). Overall, STRING identifies more disease phenotypes and a broader range of conditions, including structural cardiac disease and diverse metabolic abnormalities, reflecting its integrative network approach, than does the physical PPI network, while the physical PPI network identifies disorders that are better characterized molecularly.

One potential and interesting application of this work could be in drug design and repurposing. We identified and showed eight phenotypes (Supplementary Table 10) where drugs targeting ADR-DP proteins for some phenotypes have indications for diseases with the same phenotype. Extending this idea to other phenotypes, DREAMER can reduce the search space to find relevant protein targets for a particular phenotype. Moreover, DREAMER can be used for drug off-target prediction. Drugs with a particular ADR are expected to bind to a protein within (or close to) the identified protein sets that govern that ADR (side-effect module) (Paci et al. 2022). The reduced protein space can then be used to infer the potential off-target proteins of drugs using computational methods, e.g., Autodock and Autodock-vina (Trott and Olson 2010; Vieira and Sousa 2019), or experimental methods, e.g., based on established physicochemical methods.

The current version of our method has several limitations. DREAMER explores the mechanism of phenotypes without considering the specific variations in individual molecular profiles, which are crucial for personalized medicine. To advance our understanding in personalized medicine, one will also require access to individual-specific information. The FDA Adverse Event Reporting System (FAERS) (Center for Drug Evaluation and Research 2021) provides extensive patient information, including ADRs, drug prescriptions, dosages, and demographic details, which can be leveraged to help elucidate the mechanisms of phenotypes in the context of personalized treatments, but ultimately will require molecular-level information with which to generate individual PPIs (Maron et al. 2021).

Another limitation, yet a possible future direction, is the identification of the mechanism of phenotypes in the context of combination therapy. When drugs are approved for human use, they can be prescribed either as monotherapies or as part of combination therapies (Maciejewski et al. 2017). Combination therapy can offer synergistic benefits for treating complex or multiple health disorders, and it may result in ADRs that differ from those observed with monotherapy. For instance, in Parkinson’s disease, levodopa is prescribed to increase dopamine level and, in combination with that carbidopa is prescribed to reduce peripheral conversion reducing the ADRs such as nausea. Additionally, in Type-2 diabetes, the synergistic effects of metformin and DPP-4 inhibitors, such as sitagliptin and saxagliptin, can decrease the likelihood of ADRs by optimizing blood glucose control through distinct but complementary mechanisms. In contrast, in cancer chemotherapy, the combination of multiple medications targets cancer cells more effectively but can exacerbate ADRs, increasing risks such as severe nausea, neutropenia, infections, and significant hair loss. TWOSIDES (Tatonetti 2012) is a database that contains information on ADRs related to combinations of drugs and can be used to identify the mechanism of ADRs from the lens of combination therapy. Exploring these resources will further widen our understanding of the biological mechanisms of ADRs (Zitnik, Agrawal, and Leskovec 2018; Deac et al. 2019). In addition, certain phenotypes exhibit similarities that can also be leveraged to improve the reliability of the identification of phenotype mechanisms. Identifying similar phenotype sets can be achieved through clustering phenotypes based on the similarity in their associated drugs and diseases. The latter can be conducted using biclustering techniques, such as non-negative matrix factorization-based (Lee and Seung 2000) or knowledge graph representation learning (Hamilton 2020). In conclusion, DREAMER advances our understanding of the molecular mechanisms underlying ADRs and DPs, offering a valuable tool for improving drug safety and repurposing, thus contributing significantly to the fields of pharmacology and personalized medicine.

## Methods

### Knowledge graph construction

Our knowledge graph (KG) is composed of different node types of drugs, diseases, proteins, ADRs, and disease phenotypes (DPs). To construct this KG, we utilized data from different sources along and applied several preprocessing steps, as described in the following (Supplementary Fig. 2).

#### Drug-ADR association

To obtain the Drug-ADR association, we used the latest release of the SIDER database (SIDER 4) (Kuhn et al. 2016). SIDER is a common mono-pharmacy ADR benchmark and publicly available database that compiles information on post-marketing drug-ADR associations extracted from several public sources, including the US Food and Drug Administration (FDA). ADRs are mapped to UMLS and MedDRA terms in the SIDER database. We used the MedDRA terms to find the same DPs as ADRs. For each drug, we further removed ADRs with the same description term as its indicative disease term. Additionally, we removed ADRs linked to more than 50 drugs, as very common ADRs.

#### Disease-phenotype association

Disease-DP associations were obtained from the human phenotype ontology database (Köhler et al. 2017). DPs are termed by unique HPO identifiers. We removed DPs linked to more than 100 diseases, as very common DPs.

#### ADR-DP association

The ADR-DPs associations are collected from the Bioportal database (Rubin et al., n.d.), which reports 1,240 such relations. Among them, we preserved only ADRs associated with at least one drug that has a known protein target; and DP associated with at least one disease linked to a known gene that codes for a protein present in our dataset. In addition, in some cases, a DP (with a unique HPO id) is linked to several ADRs (with their MedDRA ids), among which we only selected one of them by random. This process resulted in a total of 649 ADR-phenotype associations.

#### Drug-target association

Drug-target associations are obtained from the DrugBank database with a premium license (Wishart et al. 2018). Drug-target associations were excluded if the drug was not included in the SIDER database.

#### Disease-gene association

Disease-gene associations were obtained from the DisGeNET database (Piñero et al. 2017). Here we used the expert-curated resource available in DisGeNET.

#### Protein-protein association

Protein-protein associations were obtained from the STRING database (von Mering et al. 2003), an extensive resource that aggregates both experimental and predicted interaction data. We only considered associations with a high confidence score (above 800) to ensure the reliability of the inferred protein interaction networks in our analysis.

Supplementary Table 1 shows a summary of node types and edge types, their number, and the databases that are used to extract this information. All edges except for STRING and physical PPI networks are obtained from the NeDRexDB platform (Sadegh et al. 2021), which is an integrative and up-to-date database for drug repurposing and disease module discovery.

### Overview of the DREAMER

To investigate the underlying mechanisms of a phenotype, DREAMER systematically identifies proteins that are highly related to both corresponding ADR and DP. For a protein to be considered relevant, it should successfully pass three statistical tests: [i] it must demonstrate significant relevance to the ADR (ADR-related protein); [ii] it must demonstrate significant relevance to the DP (DP-related protein); and [iii] there must be a significant overlap in the findings from [i] and [ii].

In particular, to obtain ADR-(or DP-) related proteins for each ADR (or DP), we first list their associated drugs (diseases). We then assign a probability to each protein based on the number of their links to the listed drugs (or diseases) as

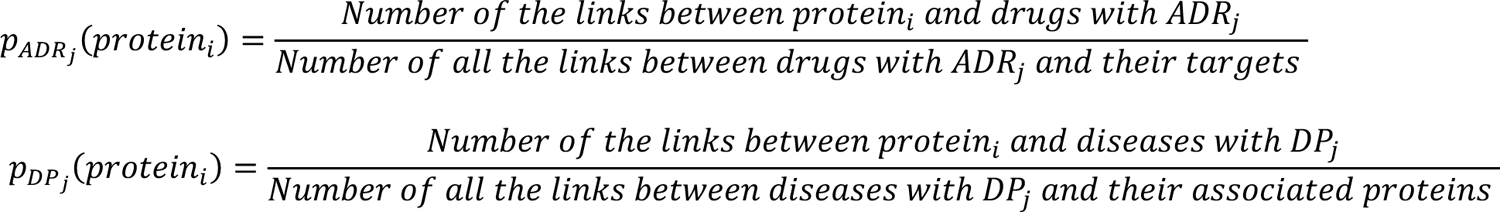

This probability vector is then used as the initial vector (seed nodes) for the diffusion algorithm (see section 4.3) over PPI resulting in two vectors of 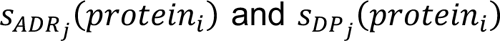, which are scores of all protein-coding genes within the interactome after diffusion. To test the significance of each protein score, we perform a permutation-based statistical test. To this end, we shuffle the protein scores of the initial vectors and obtain null diffusion scores 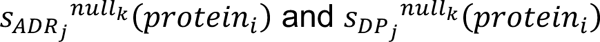 for K = 1,000 times. Lastly, we calculate a one-sided (greater) p-value for each protein score by fitting an exponential distribution on empirical null distribution and correcting it for multiple testing over all proteins in the PPI using the BH method. The final set of phenotype-related proteins is obtained from the intersection between ADR- and DP-related proteins only in case of being statistically significant in terms of hypergeometric test corrected by Benjamini-Hochberg across all phenotypes (p.value < 0.05).

### Network diffusion

Network diffusion is performed for each ADR (DP) independently through the PPI network initiated from drug targets (disease associated proteins). To this end, we use the *personalized page rank (PPR -* the R package *igraph* v1.5), which is an algorithm, developed by google to rank web pages based on their relevance to a specific user or topic (Brin and Page 1998). The PPR can capture the flow of information through the network by considering a walker that explores the network by taking steps in different directions starting from initial seed nodes. As a result, the walker visits nodes that are close to the seed nodes (probably being involved in the same mechanism) more often than the rest of the nodes in the network. The iterative formula of PPR can be defined as

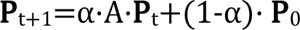

Where ***P***_t+1_ is the diffusion vector after the *t*+1th iteration, α is damping factor set to 0.7 as suggested by Hofree et al. (2013) for the STRING network. *A* is the transition probability matrix or the adjacency matrix in the PPI. Finally, ***P***_0_ is personalization vector or preference vector, which is served as the initial score.

### Significant overlap association baseline method

To assess the efficiency of the diffusion-based algorithm, we considered a baseline method also used in Kuhn et al. (2013). This method recognizes two protein sets for each phenotype, one for the ADR and the other for DP. We used the same KG (Supplementary Fig. 3), including the same ADRs and DPs for the baseline method (i.e., without the diffusion process). This method assigns a protein to an ADR (or DP) if there is a significant overlap between the set of drugs (or diseases) linked to that protein and the set of drugs (or diseases) linked to the given ADR (DP). To check for the significance of the overlap, we used a hypergeometric test. We refer to these protein sets as baseline-ADR proteins and baseline-DP proteins for ADRs and DPs, respectively.

### Significance Assessment of Overlapping Sets

To test for the significance of overlap between two sets throughout our analysis, we used the HG test. We further corrected the p-values using BH and considered α = 0.05 as the significance threshold.

## Code availability

We provide open-source R codes that allow the reader to replicate the results in this work and are available via the GitHub repository: https://github.com/faren-f/DREAMER.

## Data availability

We downloaded ADR–Phenotype, drug–ADR, drug–protein, gene–disease, gene–protein, drug– indication and phenotype–disease links from the NeDRexDB knowledgebase available at https://api.nedrex.net/. All the data are publicly available except for drug–protein links, which requires a usage license from the DrugBank dataset. The STRING PPI network was downloaded from the STRING website (https://string-db.org). The physical PPI network was downloaded from https://github.com/bwh784/PAHdrugs.

## Acknowledgements

This research was conducted as part of the DrugSiderAI project and was funded by the German Federal Ministry of Education and Research (BMBF) under grant No. 031L0306B to F.F., O.T., and J.B.; It was also supported by NIH grants U01 HG007691, R01 HL155107, R01 HL155096, and R01HL166137 to J.L.; AHA grants 957729 and 24MERIT1185447 to J.L.; EU grant HorizonHealth2021 101057619 to J.B. and J.L.; the Swiss State Secretariat for Education, Research and Innovation (SERI) under grant No. 22.00115 to J.B.; Lundbeckfonden under grant No. R347-2020-2454 to M.L.E.; the German Federal Ministry of Education and Research (BMBF) for the PoSyMed project under grant No. 031L0310A to F.F., M.L.E., Z.C., O.T., and J.B.; the European Union research under grant No. 101057619 to F.F., O.T., and J.B.; Views and opinions expressed are however those of the author(s) only and do not necessarily reflect those of the European Union or European Health and Digital Executive Agency (HADEA). Neither the European Union nor the granting authority can be held responsible for them. We thank Andreas Maier, Christiane Ehrt, and Olga Zolotareva for their assistance and valuable comments on this manuscript.

## Author contributions

F.F. developed the method and performed all the analyses; M.L.E., D.H., R.W., and J.L. provided biological interpretation; F.F. wrote the original draft of the manuscript and M.L.E., D.H., R.W., J.L. contributed to writing; F.F., M.L.E., D.H., R.W., Z.C., M.R., J.L., J.B., O.T. reviewed and edited the manuscript; F.F. and M.L.E. illustrated the figures. F.F., J.L., J.B., and O.T. conceptualized the work. O.T. and J.B. supervised the work; All authors contributed and agreed to the final version of the manuscript.

## Competing interests

The authors declare no competing interests.

## Notes

### Competing Interest Statement

The authors have declared no competing interest.

### Summary of Updates

The explanation about DataSAIL was incorrect in the previous version and has now been updated.

