## Supplementary file for "DREAMER: Exploring Common Mechanisms of Adverse Drug Reactions and Disease Phenotypes through Network-Based Analysis"

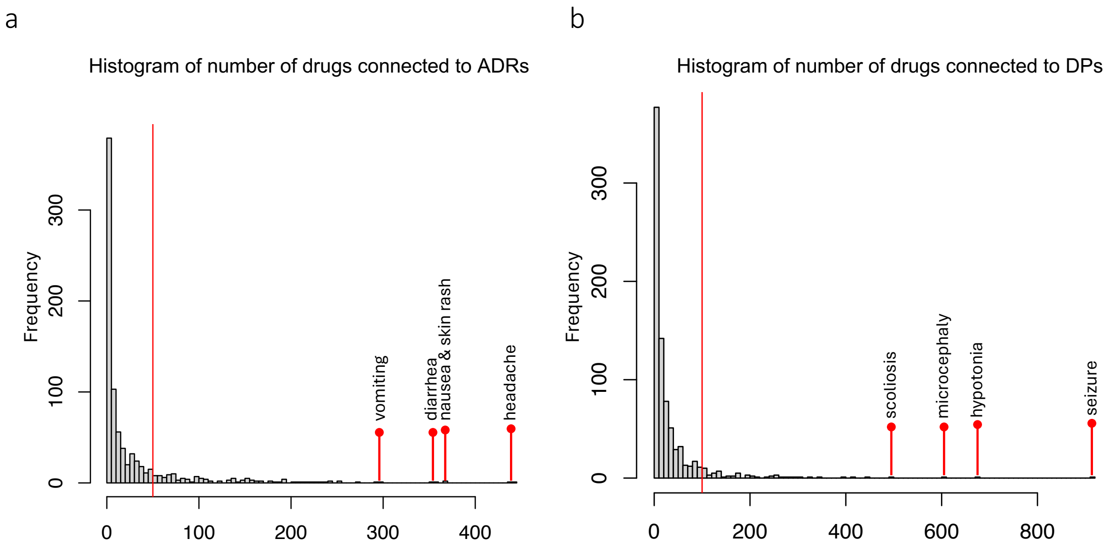


**Supplementary Fig. 1 |** The distribution of the number of A) drugs connected to ADRs, and B) diseases connected to DPs. The vertical red lines indicate the threshold used to remove ADRs and DPs that are connected to more than 50 drugs and 100 diseases respectively.

**Supplementary Table 1 | The summary statistics of nodes in our knowledge graph.**

| Node type | Initial count | Counts after preprocessing | ID type |
|---|---|---|---|
| ADR | 10368 | 649 | MedDRA |
| DP | 16874 | 649 | HPO |
| Drug | 6177 | 465 | Drugbank |
| Disease | 22825 | 2553 | Mondo Disease Ontology |
| Protein | 19385 | 12694 | UniProt |

**Supplementary Table 2 | The summary statistics of edges in our knowledge graph.**

| Edge type | Database | Initial count | # edges after preprocessing |
|---|---|---|---|

| | | | |
|---|---|---|---|
| ADR-[is the same as]-Phenotype | BioPortal | 1200 | 649 |
| ADR-[is reported for]-Drug | Sider | 217384 | 6358 |
| Drug-[has target]-Protein | DrugBank | 26537 | 5185 |
| Phenotype-[is reported for]- Disease | HPO | 227812 | 9528 |
| Gene-[associated with]-Disease | DisGeNET | 26841 | 6652 |
| Gene-[encoded by]-Protein | UniProt | 33246 | 3417 |
| Protein-[interacts with]-Protein | STRING | 11938499 | 127767 |
| Protein-[interacts with]-Protein | Physical network | – | 346535 |

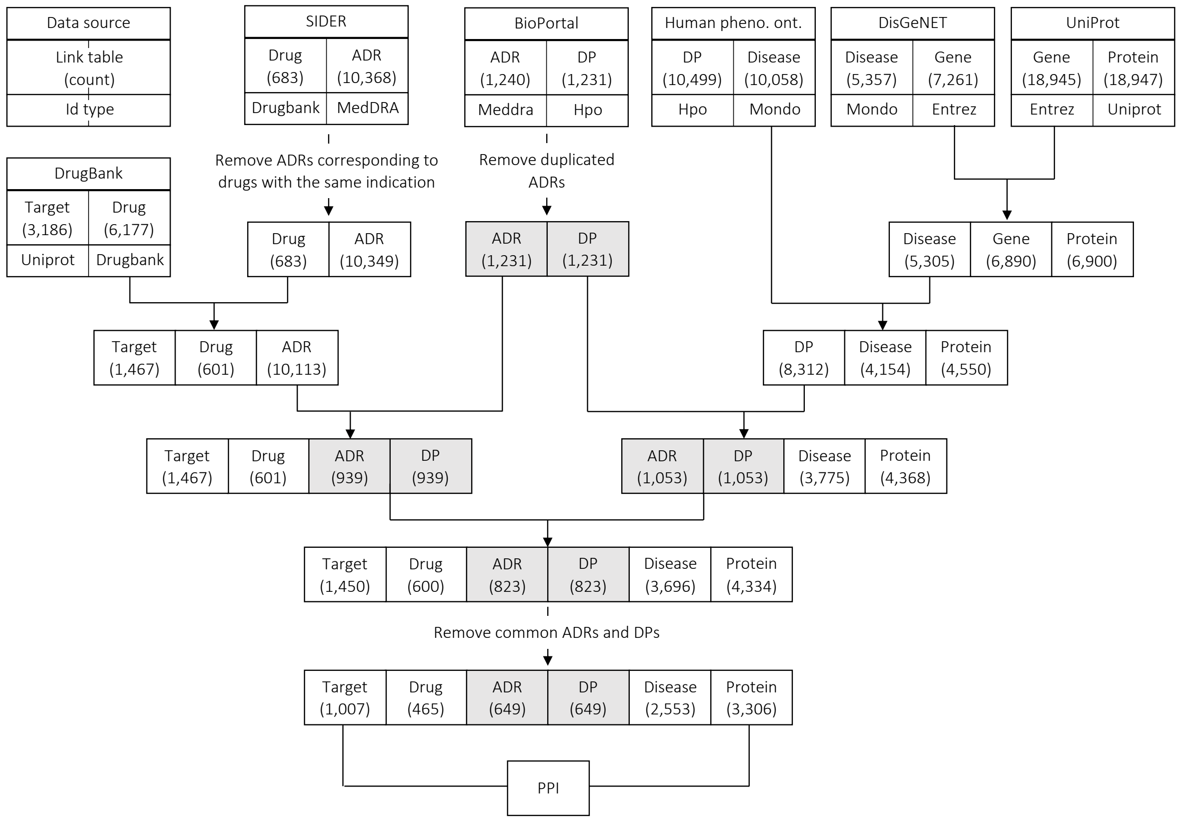


**Supplementary Fig. 2 | Data sources and pre-processing of our knowledge graph.**

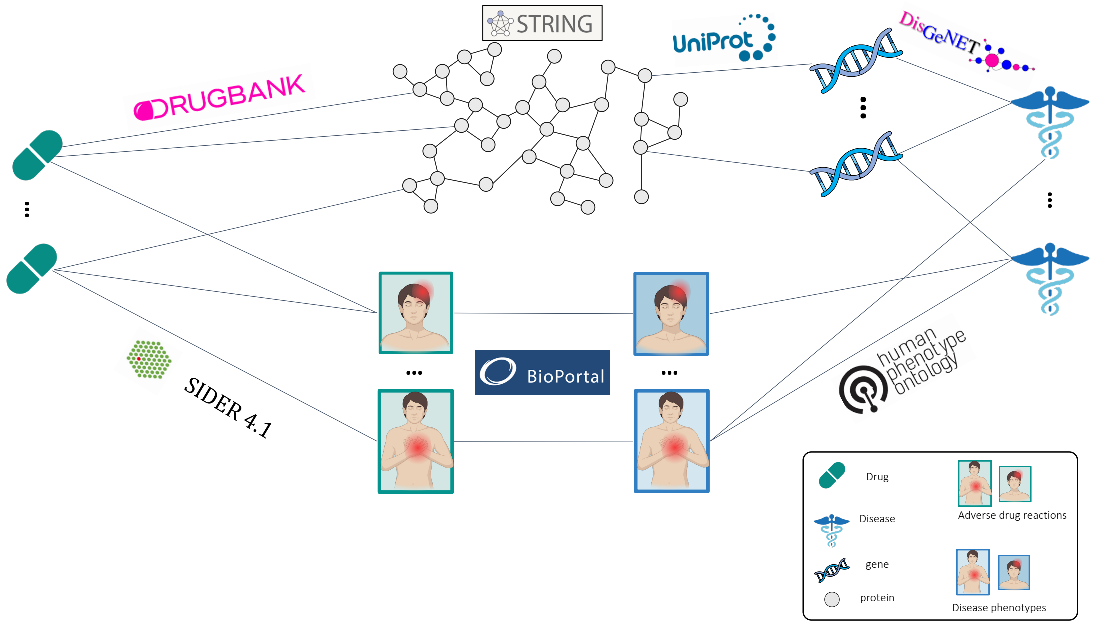


**Supplementary Fig. 3 | Our knowledge graph.**

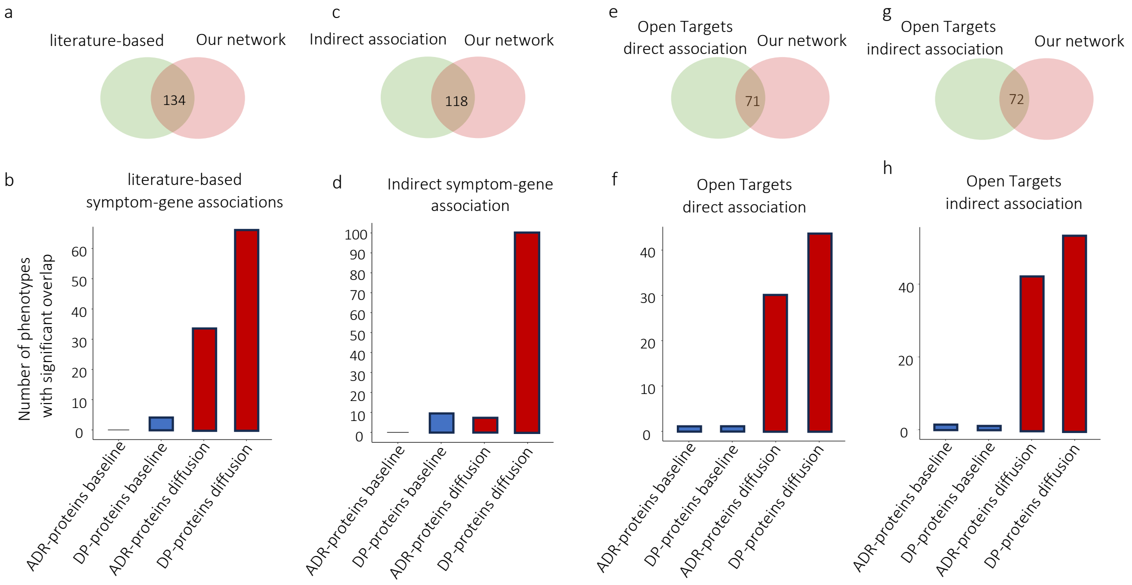


**Supplementary Fig. 4** | Reliability assessment of the identified protein set using the network diffusion algorithm on the physical PPI network; Comparison of methods using (a, b) literature-based dataset, (c, d) indirect association based on disease-phenotypes equivalent terms, (e, f) direct associations in Open Targets dataset, (g, h) indirect associations in Open Targets dataset.

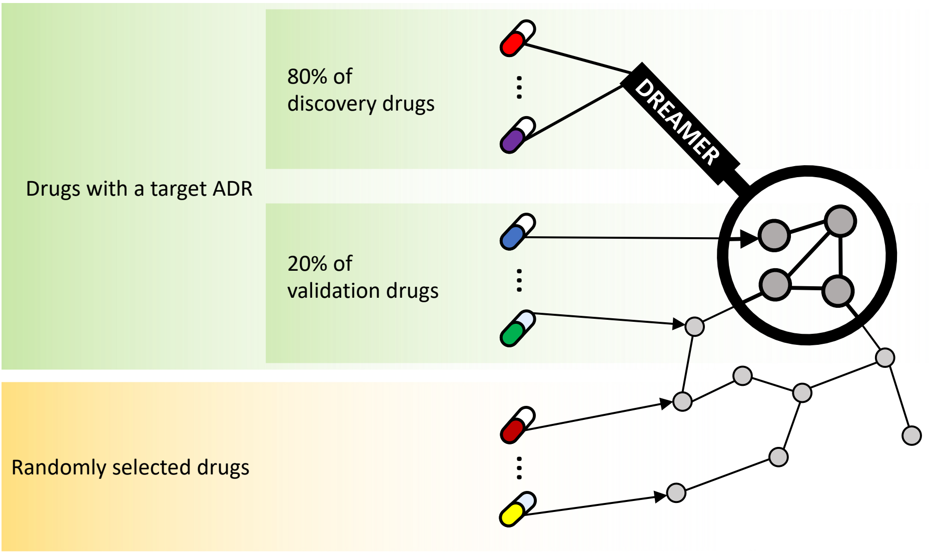


**Supplementary Fig. 5 | Schematic representing the hold-out validation analysis.** Drugs with a particular ADR are split into a discovery set (80%) and a validation set (20%). The discovery drugs are used to identify ADR-DP proteins by the DREAMER pipeline. Our analysis shows that these ADR-DP proteins are closer to the validation drugs compared to randomly selected drugs that do not have the given ADR.

**Supplementary Table 4 | The p-values obtained from Fisher's exact test after clustering of drugs based on SMILES features.**

| | Thresholds | | | | |
|---|---|---|---|---|---|
| | **X = 0** | **X = 1** | **X = 2** | **X = 3** | **X = 4** |
| **Drug holdout** | 0.003 | 0.003 | 0.9 | 0.9 | 0.9 |

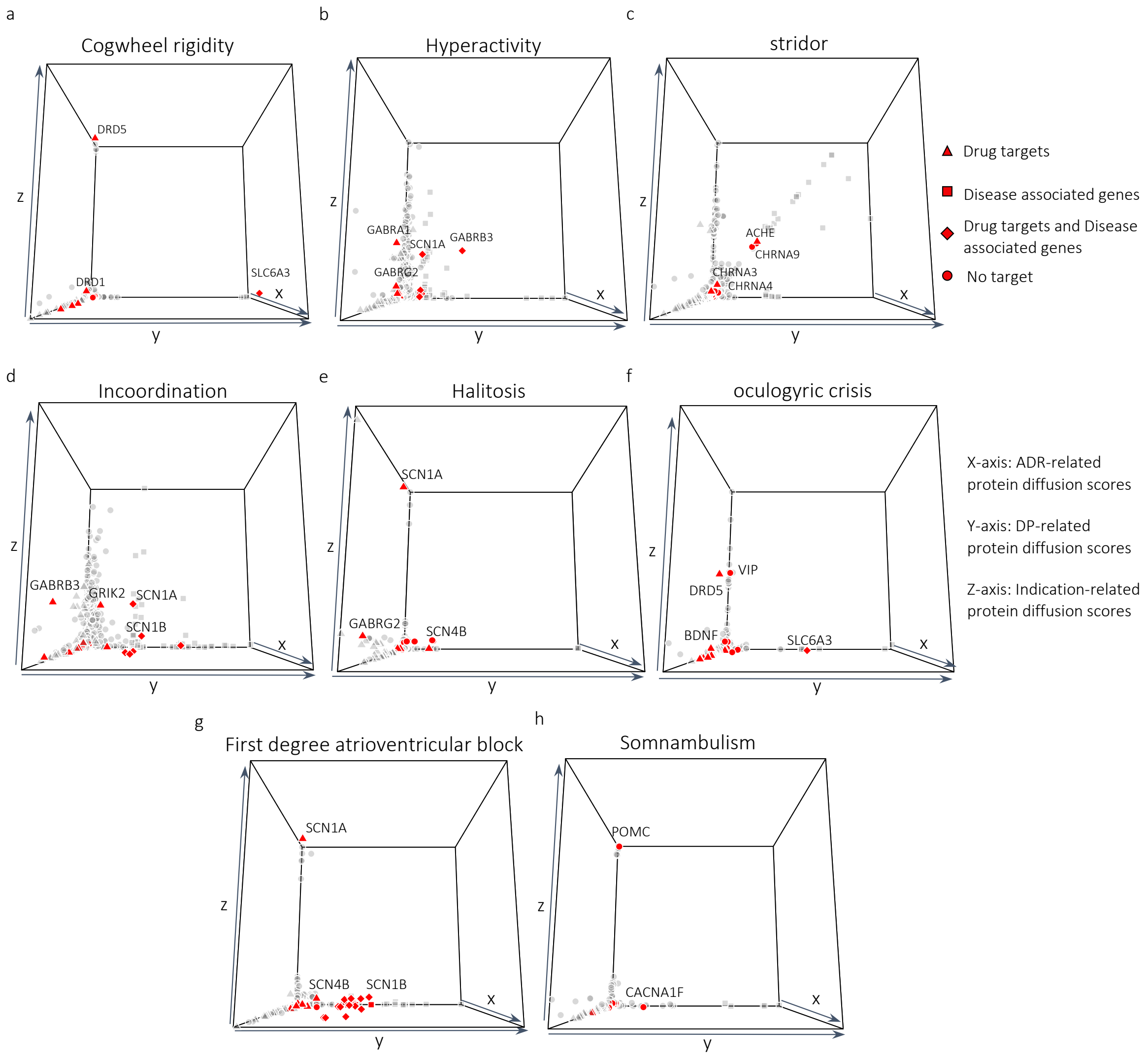


**Supplementary Fig. 6 | 3D diffusion maps.** x, y, and z-axis represent the diffusion scores of proteins from drug-targets, disease-proteins, and drug indication-proteins, respectively for (a) cogwheel rigidity, (b) hyperactivity, (c) stridor, (d) incoordination, (e) halitosis, (f) oculogyric crisis, (g) first degree atrioventricular block, (h) somnambulism. The ADR-DP proteins are indicated as red points.

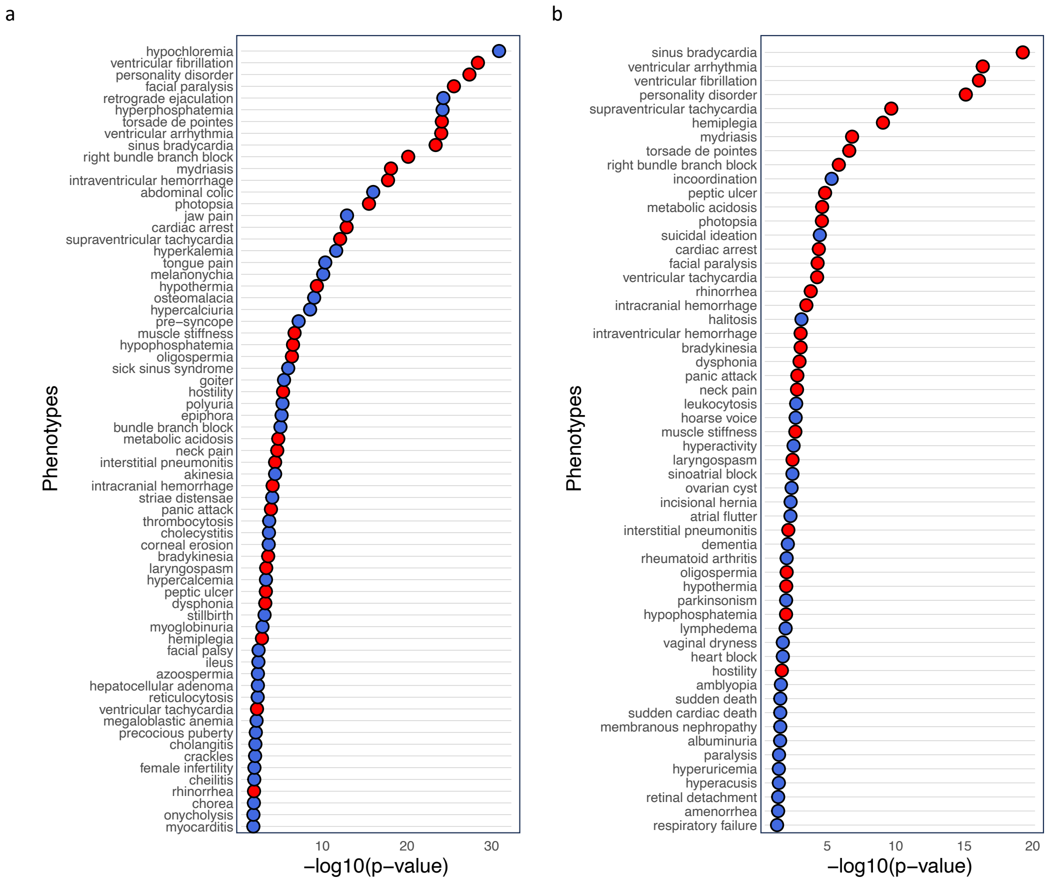


**Supplementary Fig. 7 | The ranked list of phenotypes based on the significance of their ADR-DP proteins**; applying (a) STRING network, (b) the physical PPI network. Phenotypes highlighted in "red" circles indicate those for which we identified the mechanism using an analysis based on either the STRING network or the physical PPI network.
